# *Drosophila ClC-a* is required in glia of the stem cell niche for proper neurogenic proliferation and wiring of neural circuits

**DOI:** 10.1101/482778

**Authors:** Haritz Plazaola-Sasieta, Qi Zhu, Héctor Gaitán-Peñas, Martín Rios, Raúl Estévez, Marta Morey

## Abstract

Glial cells form part of the neural stem cell niche and express a wide variety of ion channels; however, the contribution of these channels to nervous system development is poorly understood. We explored the function of the *Drosophila ClC-a* chloride channel, since its mammalian ortholog *CLCN2* is expressed in glial cells, and defective channel function results in leukodystrophies, which in humans are accompanied by cognitive impairment. We found that *ClC-a* was expressed in the niche in cortex glia, which are closely associated with neurogenic tissues. Characterization of loss-of-function *ClC-a* mutants revealed that these animals had smaller brains and widespread wiring defects. We showed that *ClC-a* is required in cortex glia for neuroepithelia and neuroblast proliferation and identified defects in a neuroblast lineage that generates guidepost glial cells essential for photoreceptor axon guidance. We propose that glia-mediated ionic homeostasis could non-autonomously affect neurogenesis, and consequently, the correct assembly of neural circuits.

## INTRODUCTION

The remarkable proliferative capacity of stem cells requires tight regulation to ensure generation of the appropriate amount of cells and tissue homeostasis during development. This regulation is controlled not only by stem cell-intrinsic programs, but also by extrinsic cues from the surrounding cellular niche. In vertebrate and invertebrate nervous systems, glia form part of the niche for neural stem/progenitor cells (Bjornsson et al., 2015; Ruddy and Morshead, 2018). In both systems, the effect of glia on neurogenic tissues has mainly been related to the secretion of factors that regulate the maintenance, proliferation, and differentiation of stem and progenitor cells.

One of the findings that changed the earlier view of glia as simply a passive structural element was the observation that glial cells expressed a wide variety of ion channels and neurotransmitter receptors (Barres, 1991; Barres et al., 1990). Although the physiological roles of several of these ion channels in glia have been described both in normal and pathological states of the mature nervous system (Black and Waxman, 2013; Nwaobi et al., 2016; Olsen et al., 2015; Pappalardo et al., 2016; Verkhrastsky and Steinhauser, 2000), the contribution of glial ion channel functions specifically in the niche during nervous system development remains poorly understood.

Among the ion channels expressed in glia, the vertebrate ClC-2 chloride channel has been proposed as one of the channels involved in K^+^ buffering, a key ionic homeostasis process in which glia are involved (Wang et al., 2017, Jentsch and Pusch, 2018). In the mature nervous system, increased neural activity leads to an increase in extracellular K^+^, which can alter neuronal excitability. To lower the concentration of K^+^, astrocytes take up the ion and distribute it to distant sites via the astrocytic syncytia. Uptake of K^+^ occurs concomitantly with uptake of Cl^−^ and water, producing transient astrocyte swelling (Bellot-Saez et al., 2017). Based on its expression in astrocytic glia, the ClC-2 channel has been proposed as one of the channels that might participate in this Cl^−^ uptake (Blanz et al., 2007; Hoegg-Beiler et al., 2014; Sirisi et al., 2017). Mutations in *CLCN2, which codes for* ClC-2, are responsible for leukoencephalopathy with ataxia (LKPAT) (Depienne et al., 2013) and ClC-2 has been related to megalencephalic leukoencephalopathy with subcortical cysts (MLC) (Hoegg-Beiler et al., 2014; Jeworutzki et al., 2012; Sirisi et al., 2017). Both conditions are characterized by vacuolization of white matter and edema, most probably as a consequence of impaired K^+^ buffering, but patients can also present learning disabilities and mild to moderate intellectual impairment. The finding that ClC-2 is expressed during development in glial precursors and is required for their differentiation (Hou et al., 2018), together with the fact that intellectual impairment can arise from connectivity defects, suggests that this channel may have additional functions during neural development. To explore this possibility, we leveraged the functional parallelisms between vertebrate and *Drosophila* glia (Chotard and Salecker, 2004; Corty and Freeman, 2013; Freeman and Doherty, 2006) and used the fly, where neurogenesis has been extensively studied, the niche is simpler than in vertebrates, and the *ClC-a* gene codes for the fly homolog of the vertebrate ClC-2 chloride channel.

The fly central nervous system contains three structures: the central brain (CB), the ventral nerve cord (VNC), and the optic lobe (OL). The CB and VNC are generated by neural stem cells called neuroblasts that delaminate from the neuroectoderm during embryonic development and give rise to larval and adult brain through two rounds of neurogenesis (Doe, 2008). The OL originates from a group of neuroepithelial cells that proliferates and separates into two crescent shaped primordia, the outer and inner proliferation centers (OPC and IPC), which produce neuroblasts and precursor cells of the different visual processing centers (Apitz and Salecker, 2014). In addition, the OL has been extensively used as a model to explore neural circuit assembly (Plazaola-Sasieta et al., 2017), primarily because the modular nature and stereotyped development of the fly eye enable easy detection of wiring defects in photoreceptors and other visual system neurons.

The cellular components in the fly niche comprise the neurogenic cells themselves (neuroepithelia and/or neuroblasts and precursor cells), the newly generated neurons, and three types of glia. Of these latter, the perineural and subperineural glia are components of the blood brain barrier (BBB) that respond to systemic nutritional cues and signal to neuroblasts to proliferate (Chell and Brand, 2010; Kanai et al., 2018; Perez-Gomez et al., 2013; Sousa-Nunes et al., 2011; Spéder and Brand, 2014). Cortex glia are large cells that lie beneath the subperineural glia. Nutritional cues and neuroblast signals alike induce cortex glia remodeling to encase neuroblasts and their immediate progeny in a chamber and older neurons individually (Read, 2018; Spéder and Brand, 2018). This close association protects neuroblasts from oxidative stress and nutritional restriction (Bailey et al., 2015; Cheng et al., 2011), and is essential for neuronal survival (Coutinho-Budd et al., 2017; Dumstrei et al., 2003; Pereanu et al., 2005; Read, 2018; Spéder and Brand, 2018). In the OL, a distinct subtype of cortex glia that expresses miRNA mir-8 (surface associated cortex glia) is in direct contact with the OPC (Morante et al., 2013). These glial cells send signals that regulate expansion of the neuroepithelium and timely transition from neuroepithelium to neuroblast (Morante et al., 2013). Connectivity is also influenced by glial cells in the visual system, where different types control photoreceptor axon pathfinding and targeting (Chotard and Salecker, 2008; Poeck et al., 2001).

The electrophysiological properties of *Drosophila* ClC-a are very similar to those of its mammalian counterpart (Flores et al., 2009; Jeworutzki et al., 2012). In addition, both ClC-2 and ClC-a are most abundant in epithelia and the brain. ClC-2 has been shown to play a role in transepithelial transport in enterocytes (Catalán et al., 2004). Similarly, *ClC-a* is also expressed in the epithelia of the fly digestive system, and is involved in transepithelial transport in stellate cells of the Malpighian tubules, the fly secretory system (Cabrero et al., 2014; Denholm et al., 2013). In the vertebrate brain, besides glia, ClC-2 is also expressed in inhibitory neurons, where it regulates neuronal excitability (Földy et al., 2010; Ratte and Prescott, 2011; Rinke et al., 2010). We were interested in the observation that *ClC-a* mRNA is expressed in glia in the embryonic fly nervous system (Kearney et al., 2004; Tomancak et al., 2007, 2002) and is highly expressed throughout development of the nervous system (Celniker et al., 2009; Rose et al., 2007), which indicates a possible role of the channel in nervous system development.

In this study, we analyzed the expression pattern of *Drosophila ClC-a* in the brain, characterized the first loss-of-function mutant alleles of this chloride channel and investigated their effects on development of the nervous system. We found that *ClC-a* is expressed in several types of glia and uncovered a role for this channel in the niche. Its expression in cortex glia, which are in close contact with OPC and IPC neuroepithelial cells and NBs, was necessary for the proper mitotic activity of these neurogenic tissues, as well as for neuron survival. One of the secondary consequences of reduced neuroblast proliferation was the significantly limited production of guidepost glial cells, which gave rise to non-autonomous neural circuit assembly phenotypes in photoreceptors. Both neurogenic and connectivity defects could be rescued by glial-specific expression of the rat ClC-2 vertebrate channel. We propose that the expression of ion channels in the glial niche can shape the development of the nervous system, controlling the number of glia and neurons generated, as well as the connectivity of the latter.

## RESULTS

### *ClC-a* is expressed in two types of glia in the developing brain: cortex glia and ensheathing glia

To characterize *ClC-a* expression in the developing brain at different larval stages (L1, L2, and L3), we used reporter lines and antibodies. One of these reporter lines expresses GAL4 under the *ClC-a* endogenous regulatory sequences (see below) and recapitulates the previously reported *ClC-a* expression pattern in Malpighian tubule stellate cells observed with an antibody against ClC-a (this study and Cabrero et al., 2014) and a ClC-a protein trap (ClC-a-GFP) (Supplementary Information and Supplementary Figure 1A-C). To visualize and monitor *ClC-a* expressing cells throughout development, we combined the *ClC-a-GAL4* line with UAS transgenes that outlined the membrane and labeled the nucleus. *ClC-a* expression was detected in L1 brains on membranes in contact with the developing OPC neuroepithelium (Figure 1A, B). Colocalization of the nuclear RFP signal with the pan glial nuclear marker Repo indicated that *ClC-a* was expressed in a subset of glial cells (Figure 1A, A’). In L2 brains, glial membranes started encasing CB neuroblasts (Figure 1C), and more membrane processes were observed deeper in the brain (Figure 1D). By late L3, the number of ClC-a^+^ glial nuclei in the CB had greatly increased and their glial membranes confined neuroblasts and their lineages in chambers (Figure 1E). A slightly deeper section showed ClC-a^+^ glial processes forming a smaller mesh (Figure 1F) which ensheathed mature neurons. In the OL, neuroblasts and progenitors were still being produced by the OPC and IPC, which continued to be surrounded by ClC-a^+^ glial membranes (Figure 1G). We also observed a glial process between the developing lamina (i.e., lamina precursor cells or LPC) and the lobula plug (LoP) (Figure 1H), establishing a boundary between these two regions, which are innervated by neurons of different origin (i.e., photoreceptors generated in the eye disc and innervating the OL through the LPC area, and distal neurons generated from the d-IPC, a region of the IPC) (Figure 1I). Similar expression patterns were observed with anti-ClC-a antibodies and the ClC-a protein trap, confirming the specificity of the *ClC-a-GAL4* driver line in the brain (Supplementary Figure 1D-I).

**Figure 1.**
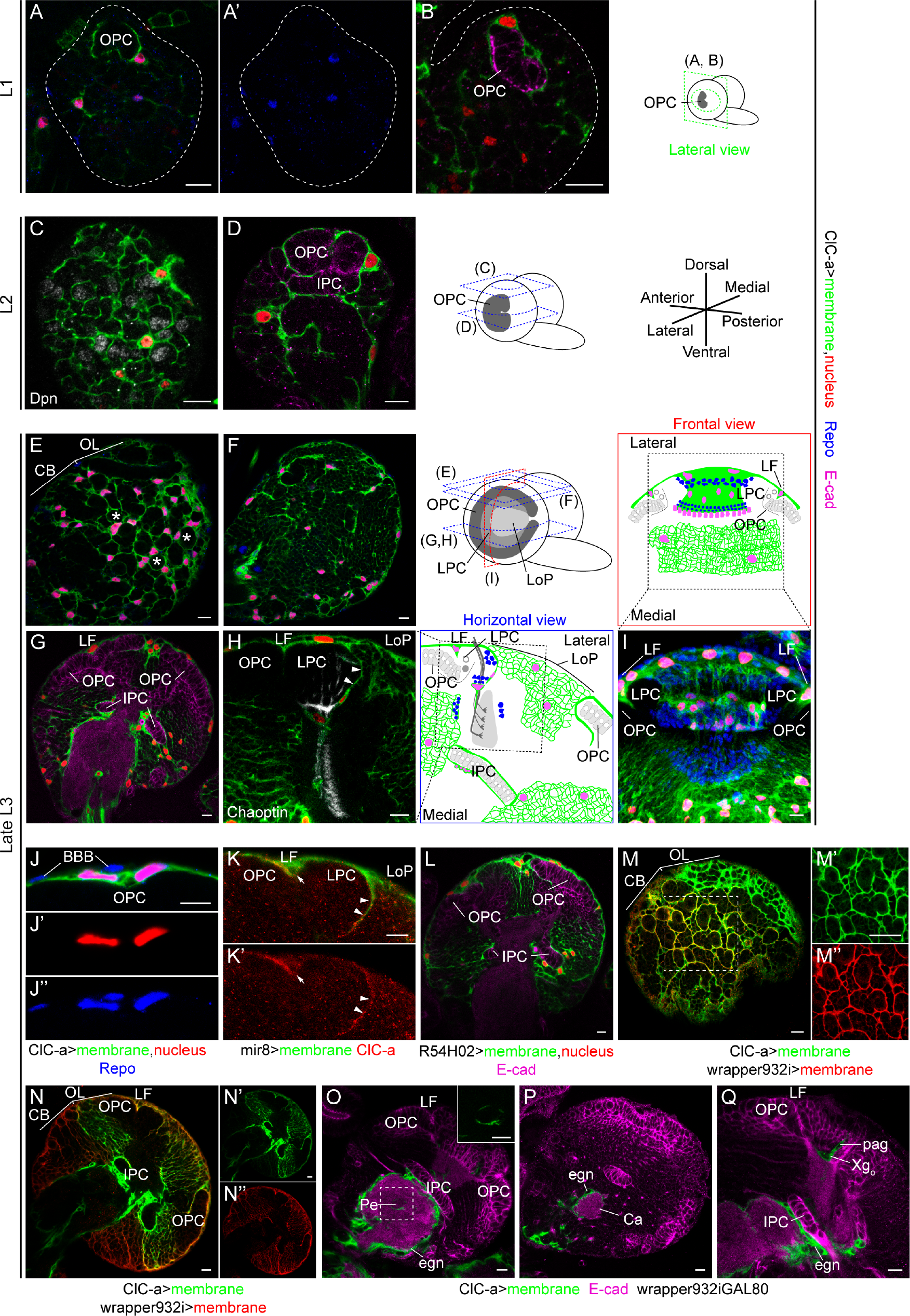
*ClC-a* is expressed in cortex glia and ensheathing glia during brain development. (A-I) Analysis of *ClC-a* expression in the developing brain. Brain illustrations show the orientation of imaging planes for the indicated panels at different larval stages. *ClC-a* specific GAL4 driver (*ClC-a-GAL4)* was used to label cellular membranes (green) and nuclei (red) of ClC-a^+^ cells. Glial nuclei were labeled with anti-Repo antibody (blue). Anti-E-cadherin (E-cad, magenta) was used to identify neuroepithelial cells. Neuroblasts and photoreceptors were labeled with anti-Deadpan (Dpn, gray) and anti-Chaoptin (24B10, gray), respectively. (A-B) Lateral view of L1 brain hemispheres outlined with a dashed line. A’ shows Repo staining from A. (C, D) Horizontal views of L2 brain hemispheres. (E, F) Horizontal views at the surface of late L3 brain hemispheres. Asterisks in E mark examples of cortex glia chambers. (G) Horizontal view through the middle of the L3 brain hemisphere. The lamina furrow (LF) is the indentation where OPC gives rise to LPCs in the lateral side. (H) Horizontal view at the same level as G, showing the region demarcated by the dashed box in the schematic on the right. Photoreceptors enter the brain through the LPC region. (I) Frontal view of a volume-rendering 3D reconstruction of the OL corresponding to the region demarcated by the dashed square in the schematic at the top. The membranes of ClC-a^+^ glial nuclei created a barrier that separated the developing lamina and the LoP. (J-Q) Identification of ClC-a^+^ glial types. (J-K) Characterization of ClC-a^+^ cells on the OPC. (J) Analysis of ClC-a^+^ nuclear position with respect to the BBB. (K) Colocalization analysis of *ClC-a* protein with *mir-8*. Expression in LF is marked by an arrow and expression between the LPC and the LoP is marked by arrowheads. (L-N) Confirmation of *ClC-a* expression in cortex glia. (L) Membrane and nuclear patterns obtained with the *wrapper* GAL4 driver (*R54H02-GAL4)* For similarity to *ClC-a-GAL4* generated patterns, compare to panel G. (M-N) Colocalization study of membrane patterns generated by the *ClC-a-GAL4* driver and a *LexA* driver version of the cortex glia *wrapper* driver (*wrapper932i-LexA*) using *UAS* and *lexAop* fluorescent reporters. (M) Horizontal view of the brain surface. To visualize *ClC-a* expression (green) in the CB (dashed region of interest), gain had to be elevated, with the consequent saturation of expression in the OL. M’ and M” show *ClC-a* (green) and *wrapper* expression (red), respectively, from the region of interest in M. (N) Deeper section into the hemisphere imaged with gain conditions to analyze OL colocalization; thus, the *ClC-a* signal (green) in the CB is very low. N’ and N” show *ClC-a* (green) and *wrapper* (red) membrane signals. (O-Q) Identification of non-cortex glia ClC-a^+^ cells as ensheathing glia. (O) *ClC-a* is expressed in neuropil-ensheathing glia (eng) surrounding CB neuropils, and tract-ensheathing glia wrapping the mushroom body peduncle (Pe, inset). (P) *ClC-a* is expressed in glia surrounding the mushroom body calyx. (Q) *ClC-a* is expressed in palisade glia and in the outer chiasm glia (Xg_o_), which wraps photoreceptor axons in their transition from the lamina to the medulla neuropils. OPC, outer proliferation center; IPC, inner proliferation center; OL, optic lobe; CB, central brain; LF, lamina furrow; LPC, lamina precursor cells; LoP, lobula plug; cxg, cortex glia; sg, satellite glia; pag, palisade glia; eg, epithelial glia; mg, marginal glia; Xg_o_, outer chiasm glia; BBB, blood brain barrier; Pe, peduncle; egn, neuropil-ensheathing glia; Ca, calyx. Scale bars represent 10 μm.

We next aimed to identify which types of glial cells expressed *ClC-a*, using cell-type-specific markers and nuclei position. We found that superficial *ClC-a*^+^ nuclei on top of the OPC neuroepithelium corresponded to a subtype of cortex glia called surface associated cortex glia (Morante et al., 2013), which lie beneath perineural and subperineural glia (Figure 1J). miRNA *mir-8* (Karres et al., 2007), a marker for this subtype of cortex glia (Morante et al., 2013), colocalized with ClC-a protein in cells covering the OPC and the process separating the LPC from the LoP (Figure 1K). Additional experiments indicated that ClC-a^+^ nuclei present on the surface of the CB and in cortical areas belonged to cortex glia. The membrane and nuclear patterns of ClC-a^+^ cells were consistent with the nuclear patterns and the membrane scaffold, also known as the trophospongium (Hoyle et al., 1986), observed with the recently described cortex glia driver *wrapper* (Coutinho-Budd et al., 2017) (compare Figure 1G with Figure 1L). In fact, there was extensive colocalization between ClC-a^+^ and wrapper^+^ membranes in the CB and OL (Figure 1M-N), including surface associated cortex glia on the OPC (Figure 1N, N”). Thus, for the sake of simplicity, we will refer to surface associated cortex glia as cortex glia.

In order to assess the presence of glial types other than cortex glia, we used an intersectional strategy whereby only ClC-a^+^/wrapper^−^ cells (i.e., non-cortex glia cells) were labeled. This revealed that *ClC-a* was also expressed in different subtypes of ensheathing glia such as neuropil-and tract-ensheathing glia. *ClC-a* was expressed in neuropil-ensheathing glia surrounding CB neuropils, including the mushroom body calyx (Figure 1O, P). For tract-ensheathing glia, *ClC-a* was expressed in glia around the mushroom body peduncle (Figure 1O), and in the OL in the outer (Xg_o_) (Figure 1Q) and inner (Xg_i_) (Supplementary Figure 2E-G) chiasm glia, cell types that wrap axonal tracts between the lamina and medulla, and the medulla and lobula complex, respectively. A detailed developmental analysis revealed expression in other glial cells in the OL, as well as in the VNC and peripheral nervous system (Supplementary Figure 2). Most of the ClC-a^+^ glial types observed in the late L3 stage persisted in the adult (Supplementary Figure 2G).

Together, these data indicate that *ClC-a* is already expressed in early development in cortex glia cells, which are in direct contact with and wrap proliferative tissues such as the neuroepithelia of the OL (OPC, IPC) and neuroblasts in the CB, forming part of the niche. *ClC-a* is also expressed in different types of ensheathing glia whose processes contribute to compartmentalization of the brain by demarcating different neuropils and neuronal tracts.

### MiMIC insertions in the *ClC-a* locus generate strong loss-of-function alleles

To explore the role of *ClC-a* in glia, we characterized a set of *Minos*-mediated integration cassette (Mi(MIC)) insertions in the *ClC-a* locus (Figure 2A). This transposon contains a gene trap cassette that leads to the formation of truncated transcripts (Venken et al., 2011) (Figure 2B). We focused on *Mi(MIC)ClC-a^05423^* and *Mi(MIC)ClC-a^14007^* alleles (from now on referred to as *05423* and *14007*), since their insertion sites were predicted to interrupt all isoforms of the *ClC-a* gene. The *ClC-a-GAL4* line we used was derived from *Mi(MIC)ClC-a^05423^* by the Gene Disruption Project (Nagarkar-Jaiswal et al., 2015a, 2015b) through recombinase-mediated cassette exchange (RMCE) replacement of the MiMIC gene trap cassette for a GAL4 cassette (Diao et al., 2015) (Figure 2B). Hence, a mutant allele is generated that expresses GAL4 under the control of *ClC-a* regulatory sequences. From now on, we will refer to it as *05423^ClC-a-GAL4^*. Initial viability characterization of these insertions over deficiency *Df(3R)PS2* (*Df*) revealed the presence of escapers (Supplementary Information). Genetic and mutant phenotype analyses (Supplementary Information and Figure 2G) indicated that allelic combinations could be ordered by strength in the following sequence: *05423^ClC-a-GAL4^/Df > 05423/Df > 14007/Df = 05423^ClC-a-GAL4^/14007 > 05423/14007 > 14007/14007*. Since it is difficult to obtain *05423^ClC-a-GAL4^/Df* or *05423/Df* animals in sufficient numbers, we mainly used *14007/Df* and *05423^ClC-a-GAL4^/14007* flies in our experiments. These two allelic combinations behave in a very similar fashion and represent a good compromise in terms of phenotypic strength and mutant animal availability. In addition, the *05423^ClC-a-GAL4^/14007* combination enables visualization in the mutant background of the glial cells that express *ClC-a* in wild type.

**Figure 2.**
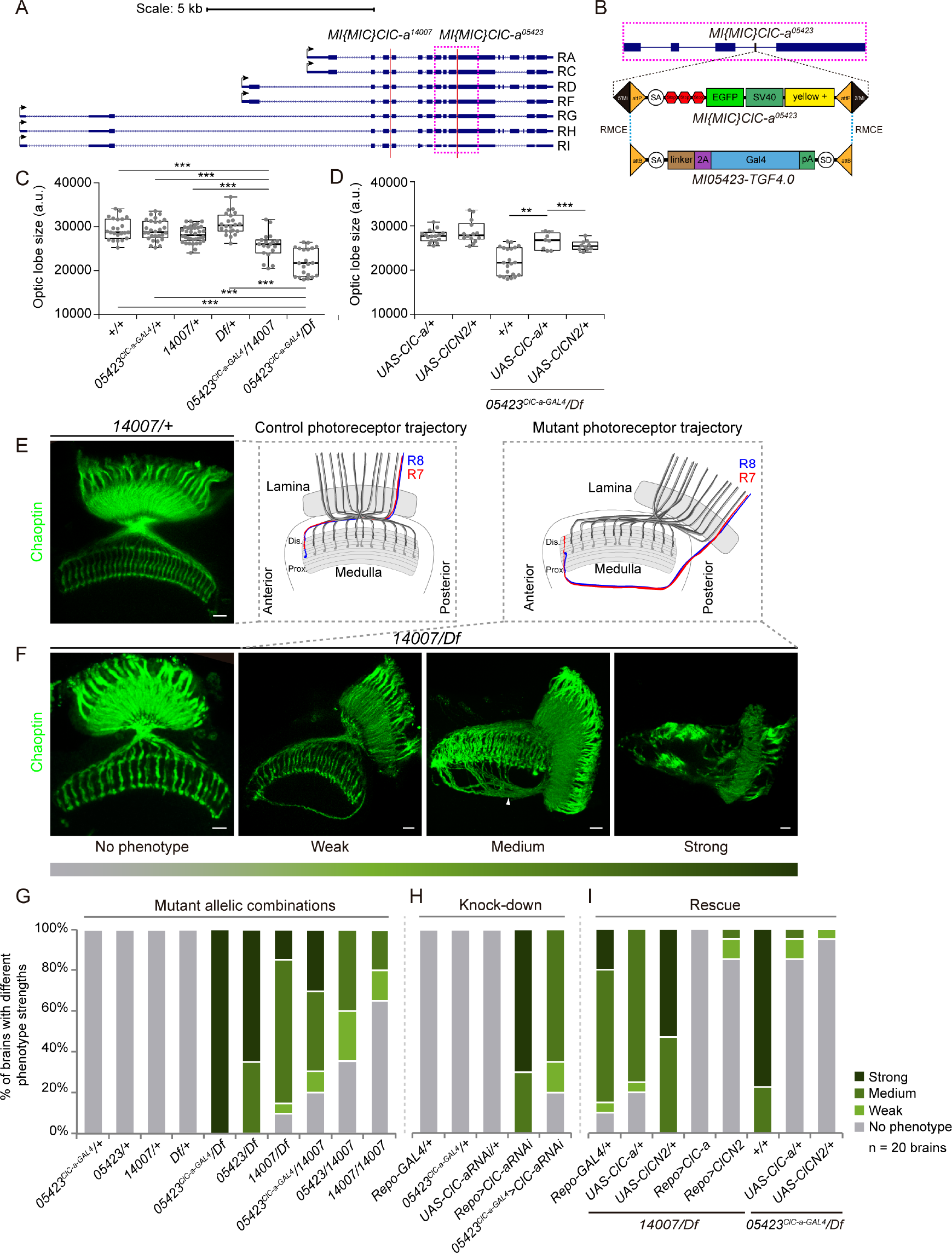
*ClC-a* mutants have smaller brains and photoreceptor guidance defects. (A) Schematic of *ClC-a* transcripts in the *ClC-a* locus and the insertion location of *Mi(MIC)ClC-a^05423^* and *Mi(MIC)ClC-a^14007^* transposons. (B) Magnification of the pink dashed box around *Mi(MIC)ClC-a^05423^* in A. The original *Mi(MIC)* transposon cassette contains a splice acceptor followed by stop codons in all reading frames, followed by the EGFP coding sequence with a polyadenylation signal. When inserted in an intron between coding exons in the orientation of gene transcription, use of the transposon’s splice acceptor generates truncated transcripts. The Trojan-GAL4 cassette swapped with RMCE to generate *05423^ClC-a-GAL4^* contains a splice acceptor that ensures the T2A-GAL4 open reading frame is included in the mRNA of the *ClC-a* gene. The T2A sequence promotes separate translation of GAL4. (C-D) Quantification of OL size in arbitrary units. *p*-values were calculated with the non-parametric Mann-Whitney test. (C) Comparison of OL size of two *ClC-a* mutant allelic combinations, *05423^ClC-a-GAL4^/14007* and *05423^ClC-a-GAL4^/Df,* and their respective controls. (D) Comparison of OL size of *05423^ClC-a-GAL4^/Df* and mutant brains in which *ClC-a (UAS-ClC-a)* or rat *CLCN2* (*UAS-CLCN2*) mRNAs were expressed in glia. (E-I) Characterization of photoreceptor guidance defects. (E) Confocal section of an adult OL of a heterozygous control animal (*14007/+*), showing the wild type photoreceptor array stained with anti-Chaoptin (24B10, green). The schematic shows the trajectory of R7 and R8 photoreceptor axons. (F) Confocal images of adult OLs from the *14007/Df* mutant allelic combination classified according to phenotype severity. For the sake of simplicity, the schematic depicts the altered trajectory of R7 and R8 axons of a single ommatidium. To show the complete trajectory of misguided photoreceptors, images for the weak and medium phenotypes are Z-projections of confocal stacks. (G-I) Photoreceptor phenotype analysis for different experiments. Phenotype penetrance and expressivity for each condition is depicted as the percentage of brains with no phenotype, weak, medium, and strong phenotypes (see Material and Methods). Heterozygous controls in (G) and (H) show no phenotype. (G) Classification of *ClC-a* mutant allelic combinations according to strength of their penetrance and expressivity. (H) Analysis of glia-specific knock down of *ClC-a* using RNAi. (I) Glia-specific rescue experiment using *ClC-a* and rat *CLCN2* mRNAs in two allelic combinations, *14007/Df* and *05423^ClC-a-GAL4^/Df*. Scale bars represent 10 μm. ** p <0.01, *** p<0.001. (See also Supplementary Figures 3 and 4)

The predicted loss-of-function nature of the MiMIC insertions characterized was confirmed by immunostaining and western blot. The *ClC-a* expression pattern observed with anti-ClC-a antibody in wild type L3 stellate cells of the Malpighian tubules and brain was not detected in any of the mutant allelic combinations tested (Supplementary Figure 3A-D). Western blots revealed that with a very low frequency, the splice machinery used the endogenous splice acceptor instead of the MiMIC one, and that there was a remnant, albeit very low, of wild type protein in mutants that was only detectable in immunoblots (Supplementary Figure 3E, F).

In summary, here we have characterized the first *ClC-a* mutants, which are strong loss-of-function alleles.

### Mutations in *ClC-a* result in smaller brains with photoreceptor guidance defects

To explore the effect of *ClC-a* mutations on brain development, we started by dissecting adult brains and searching for defects that could have a developmental origin based on *ClC-a* expression patterns in the larval brain. The observation that *ClC-a* was expressed in glia on proliferative tissues in the brain (i.e. neuroepithelia and neuroblasts) led us to hypothesize that mutant brains might be smaller than control ones, and to test this idea we measured OLs from control and mutant animals. We did indeed observe a reduction in OL size in mutants, which was particularly evident in *05423^ClC-a-GAL4^/Df*, the strongest allelic combination, and was also present in *05423^ClC-a-GAL4^/14007* (Figure 2C) and *14007/Df* (Figure 4H, I).

Given that we detected *ClC-a* expression in glial processes separating the developing lamina from the LoP and outer chiasm glial cells, we labeled photoreceptors to assess their innervation path. The compound eye of the fly is formed by some 800 units called ommatidia. Each ommatidium contains eight photoreceptors; R1-6, which terminate in the lamina forming the lamina plexus; and R7 and R8, which extend to the medulla. As rows of ommatidia form in the eye disc, photoreceptors extend axons and sequentially innervate the OL. This forms a retinotopic map and each ommatidium in the eye generates a corresponding processing unit in the lamina and the medulla neuropils. In control adult OLs (Figure 2E schematic), R-cell axons from the posterior edge of the eye enter through the posterior lamina where R1-6 stop. R7 and R8 axons traverse the outer optic chiasm and project into the anterior-most medulla; similarly, R-cell axons from the anterior region of the eye project into the posterior medulla. All R7 and R8 axons enter the medulla neuropil from its distal face and their projections align in a stereotyped array forming a retinotopic map (Figure 2E).

Analysis of *ClC-a* mutant adult OLs using a pan photoreceptor marker revealed photoreceptor guidance defects. The guidance phenotypes observed could be classified into three levels of severity based on the proportion of R-cell axons affected (Figure 2F). In brains with phenotypes classified as medium, a significant portion of posterior R-cell axons bypassed the outer chiasm, projected along the posterior edge of the medulla neuropil turning anteriorly, and extended for variable distances before innervating the medulla neuropil from its proximal face. In many cases, this resulted in posterior misplacement of the lamina neuropil. Despite the presence of these discreet bundles of misprojected axons that originate posteriorly, the photoreceptor array was maintained and mostly regular. We classified instances of few misprojected posterior axons as weak phenotypes. Strong phenotypes were characterized by severe disruption of the photoreceptor array and a posteriorly located, disorganized lamina. Despite the difficulty in identifying discreet bundles of photoreceptor axons, distal innervation was evident. These three degrees of severity showed variable penetrance and expressivity depending on the allelic combination analyzed (Figure 2G). This variability could be explained by the fact that *ClC-a* mutants were not complete nulls. A detailed analysis of mutant photoreceptors also revealed layer selection defects for R8 and R1-R6 neurons (Supplementary Figure 4A-J).

In order to confirm the requirement of *ClC-a* in glia, we performed cell-type-specific knock down and rescue experiments. In addition to the *ClC-a* driver, we also used the general glial driver Repo-GAL4 as an alternative to restrict transgene expression selectively to glia. Using these two drivers, *ClC-a* knockdown by RNAi phenocopied the photoreceptor phenotypes seen in the mutant (Figure 2H). Moreover, expression with both glial drivers of *ClC-a* and rat *CLCN2* cDNA rescued the photoreceptor phenotypes in whole mutant animals (Figure 2I). Although it has been suggested that pupal photoreceptors express *ClC-a* (Ugarte et al., 2005), we did not observe this in larval, pupal, or adult stages with the antibodies (data not shown) or reporters used in this study (Supplementary Figure 2D-G). In addition, the absence of phenotype when knocking down *ClC-a* in the eye disc or generating a full eye mutant for *ClC-a* (EGUF-hid approach, data not shown), together with the inability to rescue the guidance phenotype when expressing *ClC-a* in photoreceptors (Supplementary Figure 4K), confirmed that *ClC-a* was required in glia for photoreceptor guidance.

Remarkably, taking advantage of the *05423^ClC-a-GAL4^* allele, we observed a rescue of both OL size and photoreceptor guidance phenotypes in *05423^ClC-a-GAL4^/Df*, the strongest allelic combination, with *ClC-a* and *CLCN2* cDNA transgenes alike (Figure 2D, I). This finding indicated that brain size reduction and photoreceptor guidance phenotypes in *ClC-a* mutants are non-autonomous and dependent on chloride channel expression in glia, and that the fly and rat channels have equivalent functions.

### Expression of *ClC-a* in cortex glia is required for neuroepithelial expansion and neuroblast divisions, and is sufficient to restore brain size

In order to unravel how mutations in *ClC-a* resulted in smaller brains, we first assessed the status of glia in *ClC-a* mutants. We used the *05423^ClC-a-GAL4^/14007* allelic combination to visualize glia membranes and nuclei in the mutant background. Our analysis showed that the distribution pattern of glial cell bodies on the brain surface and deep in the cortex was similar in control and mutant animals. Although the number of nuclei/hemisphere volume ratio in the mutant was slightly reduced compared to control (Supplementary Figure 5A), importantly, the membrane scaffold appeared indistinguishable from the one observed in controls covering the whole hemisphere. As in control animals, *ClC-a* mutant cortex glia processes were in close contact with the OPC and IPC neuroepithelia (Figure 3A, B, E, F). In addition, in the OL and the CB alike, cortex glia processes formed the trophospongium. Thus, individual neuroblasts were enclosed in chambers that enlarged to adapt to their lineage expansion (Figure 3C, G), and mature neuronal cell bodies were progressively enwrapped by cortex glia processes (Figure 3D, D’, H, H’). From these observations, we conclude that mutations in the channel do not result in major morphological changes in the trophospongium formed by cortex glia.

**Figure 3.**
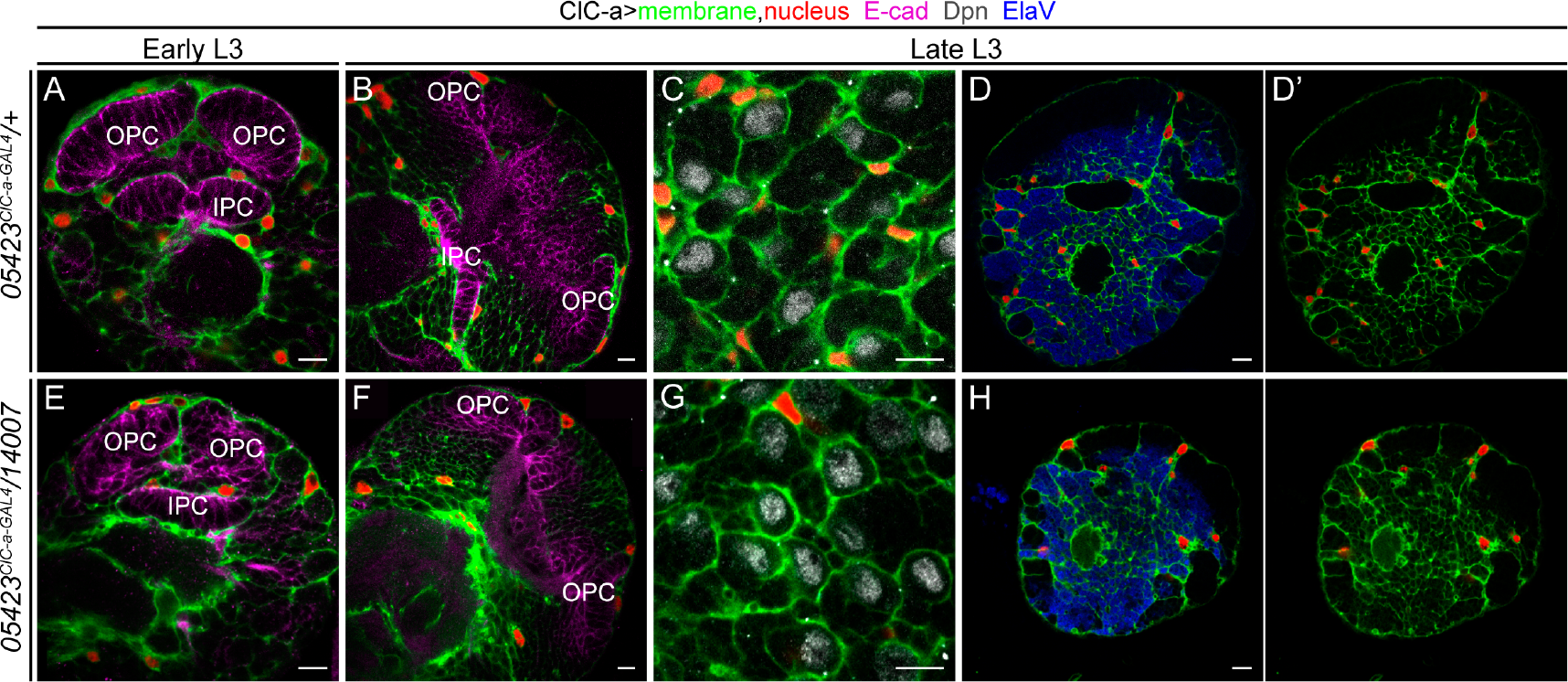
The cortex glia membrane scaffold remains unaltered in *ClC-a* mutant animals. Analysis of cortex glia membrane scaffold (green) and nuclear (red) distribution in control (*05423^ClC-a-GAL4^/+*) and mutant (*05423^ClC-a-GAL4^/14007*) brain hemispheres. Horizontal views at specified developmental times and depths are shown. (A, B) View through the middle of the early (A) and late (B) L3 hemisphere of a control animal. Anti-E-cadherin (E-cad, magenta) labels neuroepithelial cells. (C) View of the surface of a control brain stained with anti-Deadpan (Dpn, gray) to visualize neuroblasts. (D, D’) Slightly deeper view of the surface of a control brain stained with anti-Elav to visualize postmitotic neurons. (E), (F), (G) and (H, H’) panels are equivalent views and stainings in mutant animals. OPC, outer proliferation center; IPC, inner proliferation center. Scale bars represent 10 μm.

In turn, these results suggested that *ClC-a* was instead required for the proper physiology of cortex glia. Cortex glia have been shown to be essential for neurogenesis (Dumstrei et al., 2003), and since cortex glia processes are tightly associated with the OPC (surface associated cortex glia (Morante et al., 2013)) and IPC, we set out to examine whether the small OLs in mutant adult brain (Figure 2C) were a consequence of defects in these neuroepithelia. Neuroepithelia in the OL start as sheets of cells that divide symmetrically and expand until mid L3 (Ngo et al., 2010). As they do so, they bend along the dorso-ventral axis, creating a crescent shaped structure with the opening pointing posteriorly (Nassif et al., 2003). Already in late L2, while the OPC is still growing to expand the pool of prospective neuroblasts, neuroepithelium to neuroblast transition starts taking place. The lateral edge gives rise to LPC and the medial edge to neuroblasts that will produce medulla neurons and glia (Egger et al., 2007, 2010; Ngo et al., 2010; Orihara-Ono et al., 2011; Reddy et al., 2010; Wang et al., 2011a, 2011b; Weng et al., 2012; Yasugi et al., 2010, 2008). Once neuroepithelium divisions stop and the wave of differentiation continues, the OPC starts reducing in size and disappears in early pupal stages, when it is all converted into precursors and neuroblasts. A similar process takes place in the IPC, where different domains generate neuroblasts or migrating progenitors (Apitz and Salecker, 2015; Hofbauer and Campos-Ortega, 1990) until the neuroepithelium disappears.

We first checked if there were differences in neuroepithelia between control and mutant animals. For this, we stained brains with the neuroepithelium marker E-cadherin and manually segmented the tissue to generate a 3D reconstruction of these structures, which yielded information about their morphology (Figure 4A) and size (Figure 4B). In control animals in the mid L3 stage, the ends of the OPC and IPC crescents were close together. In late L3, with the addition of progeny from neuroblasts, the OL was larger and neuroepithelia crescents were wider and thinner. In comparison, in mid L3 mutant animals, neuroepithelia maintained the same crescent shape as in controls but were already clearly smaller (Figure 4B). By late L3, in most cases the OPC appeared as two separate dorsal and ventral domains with the central part absent. Similarly, part of the IPC was also missing (Figure 4A).

**Figure 4.**
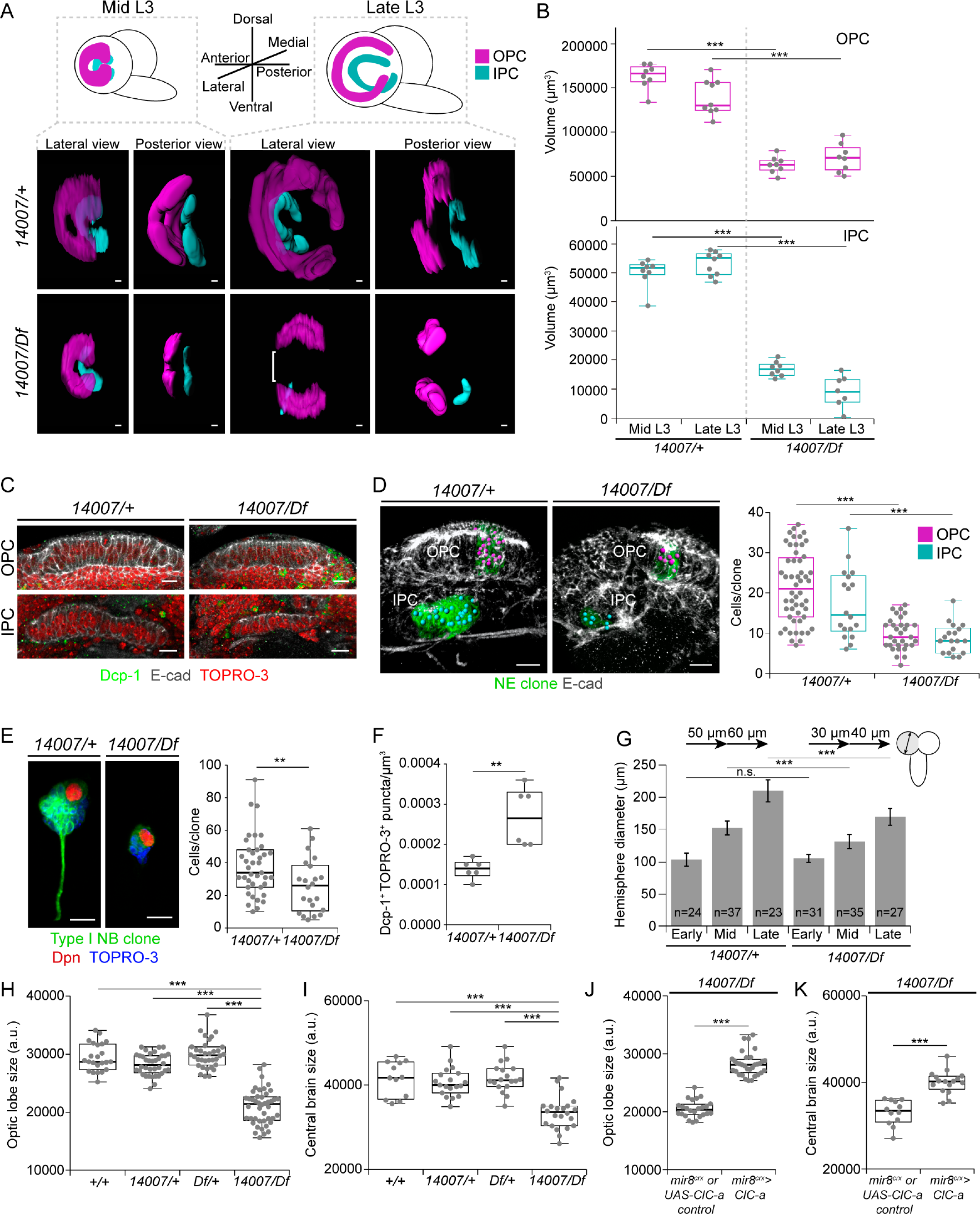
*ClC-a* is required for neuroepithelial cell and neuroblast proliferation, as well as neuronal viability, and is sufficient to rescue brain size. *p*-values of indicated comparisons were calculated with the non-parametric Mann-Whitney test unless otherwise indicated. (A) Images of surface-rendering 3D reconstructions of the OPC (magenta) and IPC (cyan) shown from lateral and posterior views, in control (*14007/+*) and mutant (*14007/Df)* brains. Bracket indicates the absence of the central domain of the OPC in mutant late L3 reconstructions. (B) Quantification and comparison of the volume in μm^3^ of reconstructed OPC (magenta) and IPC (cyan) of mid and late control and mutant animals. (C) Analysis of cell death in mid L3 OPC and IPC (E-cad, gray) of control and mutant animals using anti-Dcp-1 staining (Dcp-1, green) to label apoptotic cells. Nuclei (red) are labeled with TOPRO-3. Confocal sections show that apoptotic cells in control and mutant tissue were found outside the neuroepithelial cells. (D) Images of volume-rendering 3D reconstructions of control and mutant mid L3 OLs with mitotic clones (green) in the OPC and IPC. Anti-E-cadherin (E-cad, gray) labels neuroepithelial cells. Magenta and blue spheres represent cells in OPC and IPC clones, respectively. Quantification and comparison of the number of cells per OPC and IPC clone in the control and mutant background. (E) Images of volume-rendering 3D reconstructions of segmented mitotic clones in type I neuroblast in mid L3 control and mutant animals. The clone is labeled in green. Anti-Dpn staining (Dpn, red) identifies the neuroblast. TOPRO-3 labels the nuclei of cells in the clone. Quantification and comparison of the number of cells per clone in type I neuroblast clones in control and mutant animals. (F) Quantification and comparison of cell death (Dcp-1^+^/TOPRO-3^+^ puncta) in mid L3 brain hemispheres. (G) Graphic showing the diameter of larval hemispheres at different L3 stages in control and mutant animals. Error bars indicate standard deviation. Comparisons between control and mutant diameters at each larval stage are shown. *p*-values were calculated with the parametric Turkey’s test. The growth rate between larval stages in controls and mutants is indicated at the top of the graphic. (H, I) Quantifications and comparisons of adult OL (H) and CB (I) size for *14007/Df* animals and controls. (J, K) Quantifications and comparisons of adult OL (J) and CB (K) size in cortex glia-specific rescue experiment brains and the appropriate controls. Control brains represent genotypes for both the GAL4 driver and the UAS transgene in the mutant background since they could not be distinguished in the genetic scheme of the experiment (*mir-8^glia^* control and *UAS-ClC-a* control). For cortex glia-specific driver details, see Materials and Methods and Supplementary Figure 7. OPC, outer proliferation center; IPC, inner proliferation center. Scale bars represent 10 μm. n.s.>0.05, ** p<0.01, *** p<0.001. (See also Supplementary Figures 6 and 7)

We next wondered whether the reduction in the size of the neuroepithelial sheets was due to cell death. To test this idea, we stained larval brains with an antibody against the apoptosis marker Dcp-1 (cleaved death caspase protein-1). Although developmental cell death was taking place generally in the brain, we did not observe apoptotic cells either in control or mutant neuroepithelial cells in mid or late L3 stages (Figure 4C). Thus, the absence of cell death in this tissue suggested that defects in proliferation could be responsible for the reduction in size of the OPC and IPC at mid L3, and also for the morphological defects in late L3. In the latter, the lack of neuroepithelial cells in the OPC central domain could be due to differentiation taking place (Supplementary Figure 6) in a neuroepithelium with reduced proliferation, which would result in a premature disappearance of the tissue. To examine proliferation defects, we carried out a clonal analysis study. With this technique, once mitotic recombination has been induced in a dividing neuroepithelial cell, its progeny is labeled, and can thus be counted. Clones were generated in the L2 stage and their size was assessed 48 hours later at mid L3. Neuroepithelia clones generated in the control background (brains where cortex glia expressed *ClC-a*) presented a median size of 21 cells for OPC clones and 14.5 cells for IPC clones. Conversely, clones generated in the mutant background (brains where cortex glia did not express *ClC-a*) were significantly reduced, with a median size of 9 and 8 cells for OPC and IPC clones, respectively (Figure 4D). Thus, *ClC-a* was necessary in cortex glia for neuroepithelial expansion.

Given that *ClC-a* expressing cortex glia also cover the neuroblasts that originate from the OPC as well as the neuroblasts in the CB, we also used clonal analysis to assess neuroblast divisions in *ClC-a* mutants. The high density and proximity of neuroblasts originating from the OPC renders it difficult to obtain single neuroblast clones sufficiently apart from each other to be sure that the labeled progeny belongs to a single neuroblast. Thus, we analyzed proliferation in neuroblasts from the CB since these are sufficiently apart from each other. Importantly, both control and mutant animals showed the same number of neuroblasts; thus, CB size reduction in mutants was not due to a decrease in neuroblasts (Supplementary Figure 5B). Using the same clone induction protocol as for neuroepithelial clones, the median size of type I neuroblast clones in the control background was 34 cells, whereas the median size for clones in the mutant background was reduced to 26 cells (Figure 4E). At this same mid L3 stage, we also detected more cell death in mutant than control brains other than neuroepithelia, which were death free (Figure 4F). This result suggested that alterations in the physiology of cortex glia in *ClC-a* mutants affected the trophic role of cortex glia processes that wrap the cell bodies of the more mature neurons of the lineage (Coutinho-Budd et al., 2017; Dumstrei et al., 2003; Pereanu et al., 2005; Read, 2018; Spéder and Brand, 2018). Thus, we cannot rule out the possibility that the reduction in neuroblast clone size could be due to a combination of cell death and proliferation defects.

Together, these data suggest that the lack of *ClC-a* in cortex glia in the niche affects neuroepithelial cell and neuroblast proliferation, as well as mature neuron viability outside the niche. Consistent with both these observations, the size and growth rate of larval hemispheres was reduced in the mutant background (Figure 4G). Thus, these results are in accordance with a smaller OL (Figure 4H) and CB (Figure 4I) in adult *ClC-a* mutant brains than in those of control flies. Importantly, expression of *ClC-a* exclusively in cortex glia was sufficient to rescue the size of both structures in the adult (Figure 4J, K).

### Defects are also observed in the neuroblast lineage that gives rise to ClC-a^+^ ensheathing glia, which are necessary guideposts for photoreceptor axons innervating the medulla

In an attempt to understand how the non-autonomous photoreceptor guidance phenotype is related to *ClC-a* expression in the OL, we performed a detailed developmental expression analysis in the region where photoreceptor innervation takes place. In control L2 brains, horizontal views showed that the OPC and IPC were still juxtaposed and that ClC-a^+^ cell bodies were present on the surface of the brain and in the CB (Figure 1D). In L3 frontal views, we observed that a population of glia, which preceded the arrival of photoreceptor axons in the lamina (Dearborn, 2004; Perez and Steller, 1996), progressively positioned amid the expanding region between the OPC and IPC during the early to mid L3 stages (Figure 5A, B). This population was divided into two sets of nuclei, the ClC-a^-^ nuclei of satellite glia (Supplementary Figure 8A, B) and a population of ClC-a^+^ nuclei, with lower expression than cortex glia, which we called boundary glia (Figure 5B). From this seemingly homogenous mid L3 boundary glia population, two cell types could be distinguished in late L3 brains in frontal (Figure 5C) and horizontal views (Figure 5D, Supplementary Figure 8C): the Xg_o_ and a glial type that has never been described before. We called these latter cells palisade glia (pag) and they were positioned on the same plane as the cortex glia projection and the Xg_o_, forming a continuous glial barrier between the developing lamina and the LoP. We do not know if pag persist or which type they are in the adult (Figure 5E). Xg_o_ are considered tract-ensheathing glia, and one glial cell enwraps an average of 15 lamina-medulla fiber tracts (Kremer et al., 2017). Two independent studies have shown that Xg_o_ and Xg_i_ originate from the type II DL1 neuroblast lineage and migrate to the OL (Ren et al., 2018; Viktorin et al., 2013). We repeated DL1 lineage-tracing experiments and observed that progeny from the DL1 populated the OL following the same temporal pattern as ClC-a^+^ boundary glia (Supplementary Figure 8D-F). Hence, our data support the idea that boundary glia are DL1 progeny that differentiate into the newly described pag and Xg_o_. Quantification of boundary glia in control brains showed that their numbers increased from early to mid L3 and then dropped at late L3 (Figure 5M, Supplementary Figure 8G, H). In mutant brains, however, we observed a striking reduction in the number of boundary glial cells in mid and late L3 stages (Figure 5G-J, M). Given that no glial apoptosis was observed in the region (Supplementary Figure 8I, J), this result indicated that only very few boundary glia reached the OL in *ClC-a* mutants.

**Figure 5.**
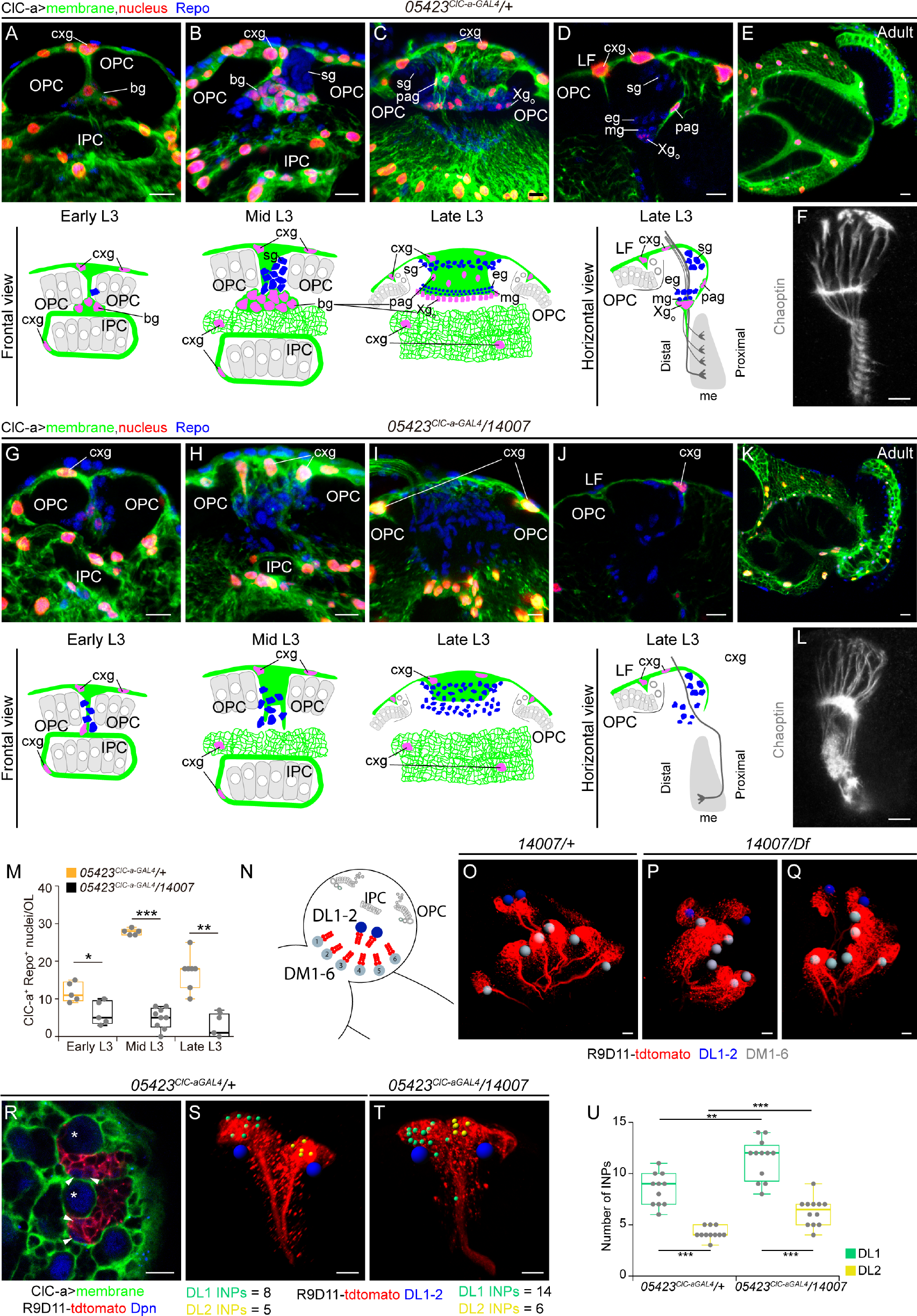
Strong reduction in a subset of ClC-a^+^ ensheathing glial cells and the neuroblast defects that caused it are observed in *ClC-a* mutants. (A-M) Developmental analysis of cells that express *ClC-a* in the OL region in control animals (*05423^ClC-a-GAL4^ /+*) and those same cells in *ClC-a* mutant animals (*05423^ClC-a-GAL4^ /14007*). Cortex glia membranes are shown in green and nuclei in red. All glial nuclei were labeled with anti-Repo antibody (blue). (A-D) Images of the ClC-a^+^ glial barrier from early to late L3 control OLs with the corresponding schematics, in frontal views (A-C) and horizontal view (D). (A-C) Volume-rendering 3D reconstructions showing the ClC-a^+^ boundary glia population in early (A) and mid (B) L3, and its division into pag and Xg_o_ in late L3 (C). (D) Confocal section. The schematic includes photoreceptors, not labeled in (D) but shown in (F). (E) *ClC-a* expression pattern in the adult OL. The inner and outer chiasms are correctly formed. (F) Photoreceptor axons (24B10, gray) in late L3 OLs. For their position relative to glia, see the horizontal view schematic. (G-J) Images showing which of the glial cells that would normally express *ClC-a* in control OLs are still present in *ClC-a* mutant OLs from early to late L3 larval stages, with the corresponding schematics, in frontal views (G-I) and horizontal view (J). (G-I) Volume-rendering 3D reconstructions. (J) Confocal section. The schematic shows the aberrant trajectory that some photoreceptor axons can take in (L). (K) *ClC-a* expression pattern in the mutant adult OL. The inner and outer chiasms are defective. (L) Photoreceptor axons (24B10, gray) in late L3 mutant OLs. For their position relative to glia, see the horizontal view schematic. (M) Quantification and comparison of ClC-a^+^/Repo^+^ nuclei in the OL region. *p*-values were calculated with the non-parametric Mann-Whitney test. (N-U) Analysis of type II DL neuroblast lineages in the CB. (N) Schematic showing the relative position of DM and DL lineages. (O-Q) Volume-rendering 3D reconstructions of late L3 control (*14007/+,* O) and mutant (*14007/Df,* P, Q) brain hemispheres showing type II lineages labeled with the R9D11-tdtomato (red). Gray and blue spheres mark the position of the DM and DL neuroblasts, respectively. (R) Confocal image showing the DL1/2 cluster lineages (red), the neuroblast (asterisk), and mature INPs (arrowheads) labeled with anti-Deadpan (Dpn, blue), and cortex glia membranes (green) surrounding the neuroblast and encasing the lineage in a glial chamber. (S, T) Volume-rendering 3D reconstructions of DL1/2 cluster lineages (red) from control (S) and mutant (T) brains where blue spheres mark the neuroblasts, smaller yellow spheres mark mature INPs of one of the lineages, and green spheres mark those from the other lineage. (U) Quantification of the number of INPs per DL lineage, showing comparisons between the number of INPs in the two lineages (green and yellow box plots) from controls and mutants. *p*-values were calculated using the non-parametric Wilcoxon matched-pairs squad rank test. Comparison of number of INPs of lineages with the highest INPs (green box plots) between control and mutants is shown. Comparison of number of INPs of lineages with the lowest INPs (yellow box plots) between control and mutants is shown. In both comparisons, *p*-values were calculated with the non-parametric Mann-Whitney test. OPC, outer proliferation center; IPC, inner proliferation center; cxg, cortex glia; bg, boundary glia; sg, satellite glia; pag, palisade glia; Xg_o_, outer chiasm glia; eg, epithelial glia; mg, marginal glia; me, medulla. Scale bars represent 10 μm. * p<0.05 ** p<0.01, *** p<0.001.

To study the cause of this marked reduction, we first used the *earmuff* R09D11 genomic enhancer-fragment driven reporter CD4-tdtomato (Han et al., 2011) to selectively label all type II neuroblast lineages and assess DL1. Type II neuroblast lineages are characterized by the generation of intermediate neural progenitors (INP) that can undergo several rounds of additional asymmetric divisions before they disappear (Boone and Doe, 2008). In control brains, there are 8 type II neuroblasts, 6 of which are positioned medially (DM1-6) and 2 laterally (DL1/2), closer to the OL (Figure 5N, O). In mutants, although we observed some brains with instances of DM mispositioning, the DL1/2 cluster was found together and laterally located with respect to the rest of the DM neuroblasts (Figure 5P, Q). However, its position with respect to the OL was sometimes changed. To assess proliferation defects in the lineage, our initial approach was to compare control to mutant DL1 clones. However, even though the clonal analysis protocol used in our study was very similar to those employed in other studies analyzing type II clones, which are identified by the presence of INPs (Dpn positive cells in the lineage), we obtained hardly any type II clones (2 out of 116 analyzed clones) and none in the DL1/2 cluster. As an alternative, we reasoned that we could use the number of INPs in a lineage as readout for proliferation (Figure 5R). Since DL1 and DL2 secondary axon tracts are extremely similar, we differentiated the two lineages through expression of *gcm-LacZ* in the DL2 lineage (Viktorin et al., 2013) (Supplementary Figure 9A), which consistently contained fewer INPs than DL1 (Supplementary Figure 9B). A comparison between control and mutant revealed that both DL1 and DL2 lineages contained a higher number of INPs in the mutant condition (Figure 5S-U). Given that we also observed proliferation defects in neuroepithelial sheets and neuroblasts, it is reasonable to suppose that one of the reasons for the marked reduction in boundary glia in mutants could be a reduced proliferation and hence accumulation of DL1 INPs. Besides proliferation defects, it is also possible that migration defects contribute to the marked reduction in boundary glia in mutant OLs. Given that the DL1/2 cluster was found at different relative positions with respect to the OL, and that the IPC, which is the region where these cells enter the OL in normal conditions, is defective in mutants, boundary glia may be hindered from reaching their final destination.

At this point, the question arises of how the marked reduction in boundary glia affects photoreceptor guidance. Since the presence of boundary glia in mid L3 coincides with the beginning of photoreceptor innervation, we next explored the spatiotemporal relationship between these two cell types in control flies. As rows of ommatidia form in the eye disc, photoreceptors extend axons that reach the OL through the optic stalk. In mid L3 stages, R8s from the first rows of ommatidia projected into the posterior part of the LPC field and their axons were located very close to boundary glia as they continued to the medulla (Figure 6A, B). Photoreceptor innervation coincided with cellular rearrangements, when boundary glia started to separate into pag and Xg_o_ glia. Thus, in slightly older brains, R1-6 axons stopped and formed the lamina plexus above the boundary glia cells that would become Xg_o_, and R8 axons traversed the outer optic chiasm, passing very close to the Xg_o_ (Figure 6C, D) and continued to the medulla, innervating it through its distal face (Figure 5F). Hence, photoreceptors are in close proximity to pag and Xg_o._ Conversely, in mutant brains, the marked reduction in boundary glia, and consequently in Xg_o_, caused posterior R8 axons to skip the outer chiasm and innervate the medulla from its proximal face (Figure 5L). The severity of initial photoreceptor guidance errors determined the strength of the adult guidance phenotypes. Consistent with Xg_o_ and Xg_i_ originating from DL1, in *ClC-a* mutants chiasms did not properly form, which resulted in altered positioning of OL neuropils in the adult brain (compare Figure 5E with Figure 5K).

**Figure 6.**
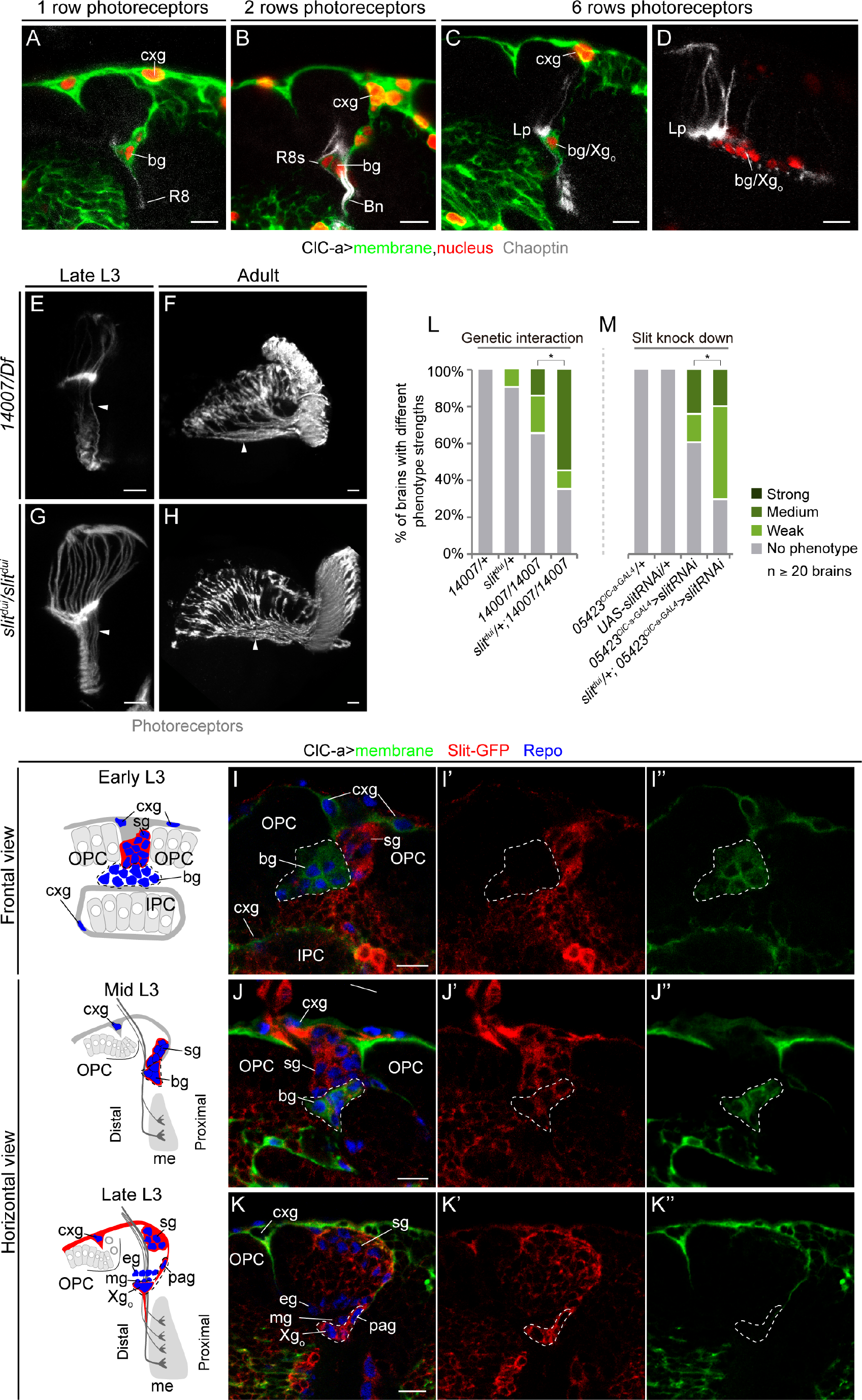
Boundary glia, which express the chemorepellent molecule Slit, are in close contact with photoreceptor axons as they innervate the OL. (A-D) Spatiotemporal relationship between photoreceptors and boundary glial cells. Number of photoreceptor rows was inferred from 24B10^+^ rows in the eye imaginal disc. (A, B) Horizontal views of mid L3 optic lobe showing ClC-a^+^ glia and one (A) and two (B) rows of R8 photoreceptors (Chaoptin, gray). (C) Same view and staining as panels A and B of a slightly older brain innervated by six rows of photoreceptors. (D) Frontal view showing transversal sections between the line of Xg_o_ cell bodies of photoreceptors on their way to the medulla. (E, F) Larval (E) and adult (F) examples of photoreceptor (24B10, gray) phenotypes in *ClC-a* mutants classified as strong. (G, H) Larval (G) and adult (H) photoreceptor (GMR-GFP, gray) phenotypes in *slit^dui^* mutants. Arrowheads show misguided axons innervating the medulla from its proximal face. (I-K) Developmental analysis of Slit expression in glial cells in the barrier. Schematics for the view in each of the stages analyzed are shown. (J) and (K) schematics include photoreceptors for orientation although they are not labeled in the images. Anti-Repo (blue) was used to label glial nuclei. A Slit-GFP protein trap (*sli[MI03825-GFSTF.2]*) that outlines membranes of *slit* expressing cells (Supplementary Figure 10) was used to visualize the *slit* expression pattern (red, I’-K’). ClC-a^+^ boundary glia (green, I”-K”) are outlined (white dashed line) in (I-I”,J-J”). Xg_o_ and palisade glia are outlined in (K-K”). Although *ClC-a* expression is downregulated in (K”), we have shown that they express *ClC-a* in other panels (Fig 1H, Fig 5D, 5P, Supplementary Figure 6K). (I) Frontal view of an early L3 OL. (J) Horizontal view of a mid L3 OL. (K) Horizontal view of a late L3 OL. (L, M) Phenotype analysis for *slit/ClC-a* genetic interaction (M) and *slit* knockdown (L). Phenotype penetrance and expressivity for each condition is depicted as the percentage of brains with no phenotype, weak, medium, and strong phenotypes. N ≥ 20 brains for each condition. *p*-values were calculated with the Chi square test. cxg, cortex glia; bg, boundary glia; Bn, Bolwigs nerve; Lp, lamina plexus; Xg_o_, outer chiasm glia; sg, satellite glia; OPC, outer proliferation center; IPC, inner proliferation center; eg, epithelial glia; mg, marginal glia; pag, palisade glia; me, medulla. Scale bars represent 10 μm. * p<0.05.

Developmental guidance defects and adult outcomes of *ClC-a* mutants are both extremely similar to OL specific *slit* mutants (Figure 6E-H) and *robo3* mutants (Pappu et al., 2011; Tayler et al., 2004). The secreted chemorepellent molecule Slit and the Robo family of receptors (Robo, Robo2, Robo3) have been implicated in preventing photoreceptor axons from mixing with distal neuron axons from the LoP during development, hence maintaining compartmentalization of this region of the developing brain (Tayler et al., 2004). While receptors have been shown to be required in neurons, *slit* reporters suggest that Slit protein in the region could be contributed by Xg_o_ (Pappu et al., 2011; Tayler et al., 2004). A detailed developmental analysis of glial barrier assembly allowed us to unequivocally characterize the temporal and cellular expression pattern of *slit* with respect to photoreceptor innervation. To this end, we characterized and used a MiMIC-based protein trap line for Slit (Supplementary Information, Supplementary Figure 10). Our analysis indicated that Slit was already being expressed in boundary glia in mid L3 (Figure 6I-K), when photoreceptors innervate the brain and their axons come into close proximity with these glial cells. Moreover, removal of one copy of *slit* enhanced the *ClC-a* photoreceptor guidance phenotype, suggesting a genetic interaction between these two genes (Figure 6L), while knocking down *slit* in ClC-a^+^ glia in the barrier recapitulated photoreceptor guidance defects (Figure 6M).

Based on our present results and previously published studies (Fan et al., 2005; Pappu et al., 2011; Suzuki et al., 2016; Tayler et al., 2004), we propose that the substantial reduction in boundary glia is most probably due to a combination of proliferation and migration defects, which results in a significant reduction in Slit protein in the region. As a consequence, photoreceptors that innervate the OL close to the glial boundary fasciculate with the axons of neurons in the LoP known to innervate the medulla from its proximal site.

### Expression of *ClC-a* exclusively in cortex glia is sufficient to restore ensheathing glia guidepost cells and rescue photoreceptor guidance defects

To test whether *ClC-a* expression in cortex glia was sufficient to regulate DL1 proliferation, and assess if *ClC-a* expression in boundary glia (cell type classified as ensheathing glia) played any role in photoreceptor guidance, we performed a cell-type-specific rescue experiment. Because no reporter has yet been described that exclusively labels boundary glia before photoreceptor innervation, we carried out a cortex glia-specific rescue. We reasoned that with a cortex glia-specific driver, we could rescue the generation of boundary glia from DL1 and at the same time avoid *ClC-a* expression in boundary glia (Figure 7A). Since we could not specifically label boundary glia in this experiment, we used Repo to mark and count glial nuclei in the region in mid L3, when the first photoreceptors begin to innervate the brain. At this time, the glial population is compact and easy to identify, whereas in late L3, additional ClC-a^-^ glia such as epithelial and marginal glia appear in high numbers and complicate counting. In control animals, mid L3 glia nuclei included ClC-a^-^ satellite glia and boundary glia (Figure 7A, B). In mutants, the number of glial cells was reduced to half due to the marked reduction in boundary glial cells (Figure 7A, B), but expression of *ClC-a* exclusively in cortex glia resulted in an almost complete rescue in the number of glial cells present in the barrier region in mid L3 (Figure 7A, B). More importantly, this boundary glia rescue also rescued the photoreceptor guidance phenotype (Figure 7C). Surprisingly, autonomous *ClC-a* expression in boundary glia was not necessary for their viability, for migration from their point of origin in the CB to position themselves in the OL, or for Slit secretion, since photoreceptor guidance defects were fully rescued when boundary glia were in their position but did not express *ClC-a.* Thus, we conclude that the strong reduction in boundary glia and the photoreceptor guidance phenotypes are a secondary consequence of the *ClC-a* requirement in cortex glia and its function in neuroblast proliferation.

**Figure 7.**
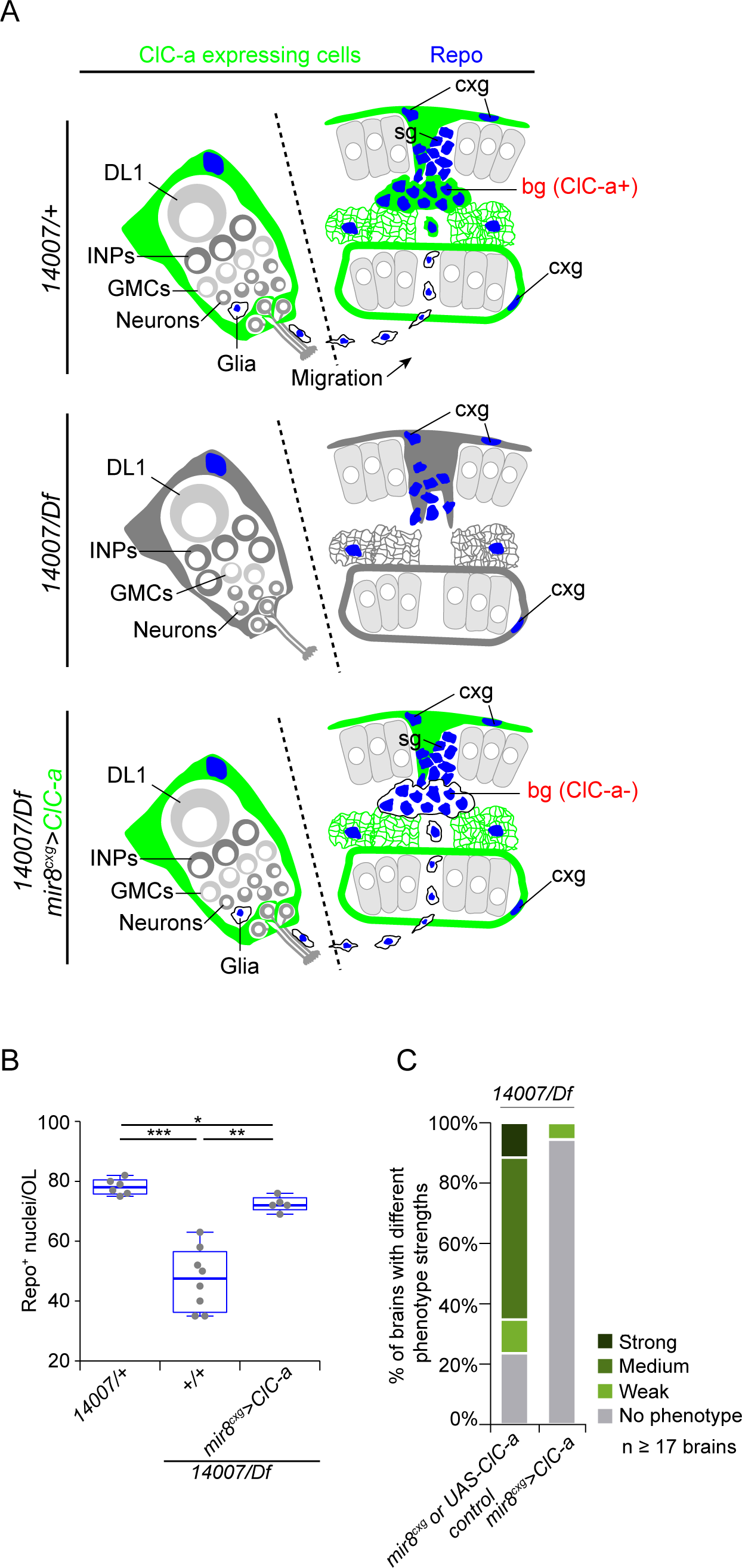
*ClC-a* expression exclusively in cortex glia rescues the formation of boundary glia and photoreceptor guidance defects. (A) Schematics depicting the cortex glia-specific rescue experiment. Frontal views of mid L3 OLs and the DL1 lineage: control (*14007/+*) showing glial nuclei in blue (Repo) and *ClC-a* expression in green in the CB in cortex glia surrounding the DL1 neuroblast and its progeny, and in the OL in cortex glia on the neuroepithelia and boundary glia; a *ClC-a* mutant (*14007/Df*) showing the absence of *ClC-a* expression and boundary glia; and an animal where *ClC-a* expression has been exclusively restored in cortex glia (*mir-8^cxg^*), resulting in the recovery of boundary cortex glia that do not express *ClC-a* because they are a subtype of ensheathing glia. (B) Quantification and comparisons of glial nuclei (Repo, blue) in control, mutant, and rescue animals. *p*-values were calculated with the non-parametric Mann-Whitney test. (C) Quantification of photoreceptor guidance phenotypes in control and rescue brains. Control brains represent genotypes for both the GAL4 driver and the UAS transgene since they could not be distinguished in the genetic scheme of the experiment (*mir-8^glia^* control and *UAS-ClC-a* Scale bars represent 10 μm. * p<0.05, ** p<0.01, *** p<0.001.

### Mutations in *ClC-a* result in widespread wiring defects

Although we have characterized the origin of the guidance defects seen in photoreceptors, wiring defects are not restricted to this cell type. The position and morphology of neuropils in the visual system of *ClC-a* animals indicate that the wiring of many other neurons in this system is also probably affected (compare Figure 5E to K). Moreover, we also observed defects in CB structures such as mushroom bodies (MBs). Each hemisphere contains one MB, which is formed by the neurons derived from four special type I neuroblasts that never enter quiescence. These neurons extend dendrites forming the calyx, and axons project into a fascicle called the peduncle that splits into two branches called lobes (Figure 8A). Similar to photoreceptors, mushroom bodies are neural structures that are highly dependent on glia-neuron interactions. It has been shown that glia wrap the peduncle and the lobes during development (Spindler et al., 2009) and in the adult (Kremer et al., 2017), and that different type II DM neuroblasts contribute glia that associate with the mushroom body (Ren et al., 2018). In control animals, ClC-a^+^ glia surrounded the MB calyx (Figure 8B) and the peduncle (Figure 8D). Newly differentiated, FasII^−^ neurons projected their axons through the center of the peduncle, generating a ring-like FasII^+^ pattern labeling the oldest neurons (Figure 8C). In *ClC-a* mutant animals, axons often misprojected into the calyx (Figure 8F) and FasII staining filled the center of the peduncle, suggesting that newly generated axons did not project through the center of this structure (Figure 8G). In addition, the peduncle was much thinner (Figure 8G), although it seemed that ClC-a^+^ glia continued to surround it. Comparison of control and mutant brains stained with antibody against N-cadherin, which labels neuropils, revealed that the calyx, which in controls appeared deep in the brain (Figure 8E), was more superficial in mutants (Figure 8I). MB clones (Figure 8J) confirmed defects in the calyx and the peduncle (compare Figure 8K-M and O). In MB clones in the control background, axons from the clone stayed together in a bundle and extended into the center of the peduncle (Figure 8L). In instances where MB clones in the mutant background extended axons into the peduncle (Figure 8M), these axons defasciculated and projected into the peduncle through its periphery, leaving older axons in the center (Figure 8N). In clones with strong phenotypes, almost all axons terminated in the calyx and the peduncle was barely visible (Figure 8M). Interestingly, these defects are very similar to those observed when cortex glia and neuropil glia are eliminated: abnormal mushroom body morphologies including splaying of axons and misguidance, and a misshapen superficial calyx due to premature fusion of the four MB lineages in the cortical region (Spindler et al., 2009). Thus, as observed for photoreceptor guidance phenotypes, MB defects in *ClC-a* mutants may be due to reduced production of glia associated with MB circuitry, whether that glia is ClC-a^+^ or not. In summary, since guidance defects in the *ClC-a* mutant seem to be widespread, we propose that the *ClC-a* requirement for proper circuit assembly is not restricted to the OL but is general to the brain.

**Figure 8.**
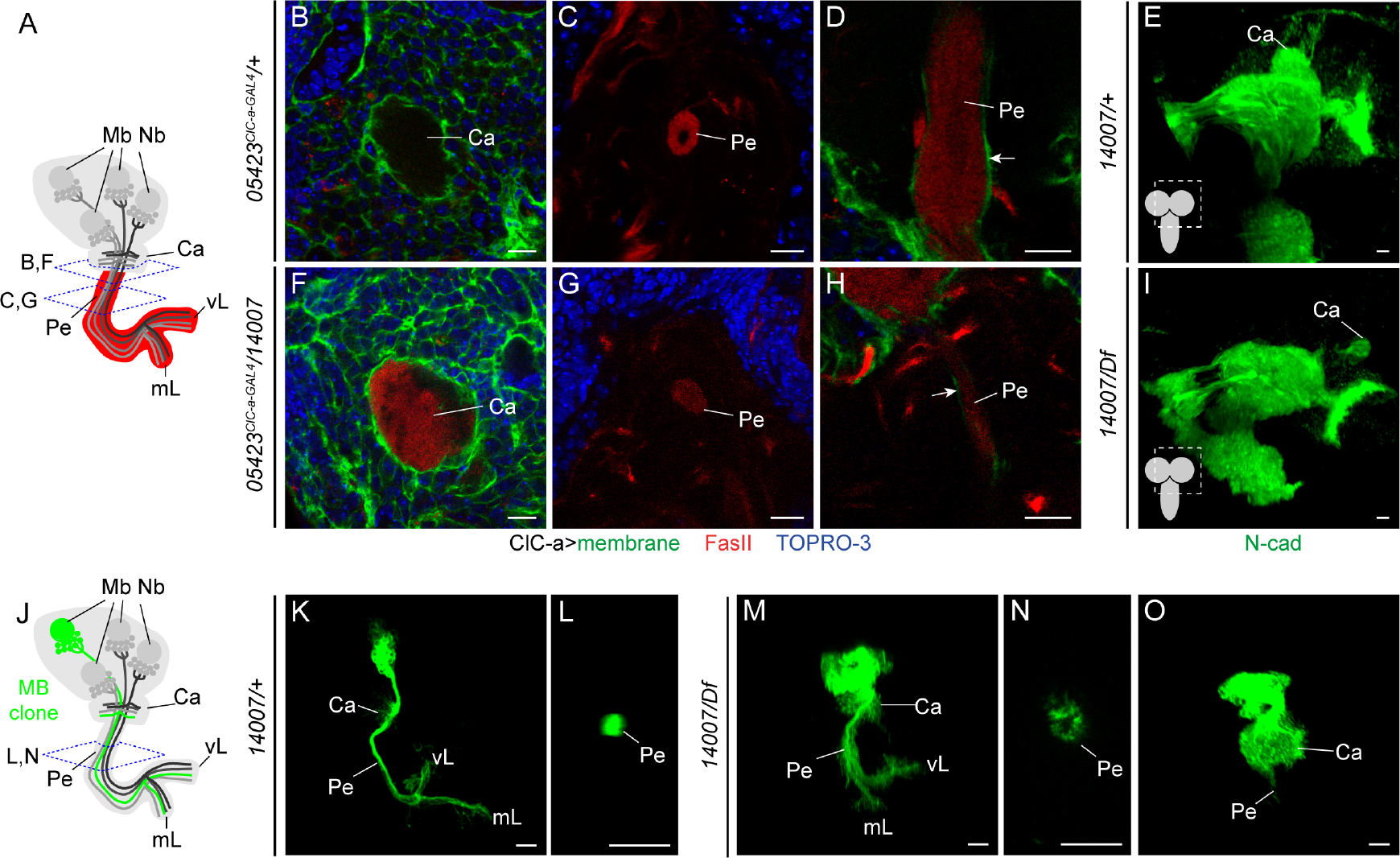
Guidance defects in mushroom body neurons. (A) Schematic of a mushroom body (MB) in one hemisphere. Dashed lines indicate the position of imaging planes and associated letters indicate correspondence to panels. The axonal component of the MB, which consists of the peduncle and lobes, is shown in red, representing anti-Fasciclin II (FasII) antibody staining. (B-E) Mushroom body analysis in control brains (late L3 *05423^ClC-a-GAL4^ /+* or mid L3 *14007/+*). (B) Confocal section though the calyx region of a control brain showing ClC-a^+^ glial membranes (green) and all nuclei (blue, TOPRO-3). (C) Transversal section through the peduncle of a control brain. (D) Longitudinal section of the peduncle showing ClC-a^+^ tract-ensheathing glia surrounding it (arrow). (E) Volume-rendering 3D reconstruction of a control brain hemisphere showing N-cadherin positive neuropils. (F-I) Mushroom body analysis in *ClC-a* mutant brains (late L3 *05423^ClC-a-GAL4^ /14007* or mid L3 *14007/Df*) with the same staining as the equivalent control panels. Compare to panels (F) to (B), (G) to (C), (H) to (D), and (I) to (E). (J-O) Schematic (J) and volume-rendering 3D reconstructions and confocal sections of mushroom body neuroblast clones in control (K, L) and *ClC-a* mutant (M-O) brains labeled in green. (K) Control clone. (L) Cross section of a control clone at the level of the peduncle. (M) Mutant clone. (N) Cross section at the level of the peduncle of a mutant clone in M. (O) Mutant clone with a strong phenotype. Ca, calyx; Pe, peduncle; mL, medial lobe; vL, vertical lobe. Scale bars represent 10 μm.

## DISCUSSION

In this study, we have shown that the *ClC-a* chloride channel function in the glial niche has a non-autonomous but profound effect on two key aspects of neural development: the generation of neurons and glia in the appropriate numbers, time, and place through its role in regulating the proliferation of neurogenic tissues, and in consequence, the correct assembly of neural circuits. Importantly, the fact that the fly (ClC-a) and rat (CLC-2) chloride channels rescue brain size and guidance defects suggests that both can perform the same physiological function. Concomitant defects in neuroblast proliferation and photoreceptor targeting have been observed in other studies (González et al., 1989; Kanai et al., 2018; Zhu et al., 2008), and it has been proposed that the

Activin signaling pathway is required to produce the proper number of neurons to enable proper connection of incoming photoreceptor axons to their targets (Zhu et al., 2008). Interestingly, mutations in the proneural gene *asense*, which is expressed in type I neuroblasts and INPs, has adult targeting phenotypes that are extremely similar to the ones observed in *ClC-a* mutants (González et al., 1989). Along the same lines, our study links *ClC-a* photoreceptor guidance phenotypes to INP proliferation defects, and furthermore identifies the INP-derived cellular population required for proper photoreceptor axon guidance.

In addition to leukoencephalopathy, patients with mutations in *CLCN2* or altered function of the channel also show cognitive impairment. Similarly *CLCN2* mutant mice develop widespread vacuolization that progresses with age, but besides photoreceptor and male germ cell degeneration, they do not display immediately visible behavioral defects (Blanz et al., 2007; Bösl et al., 2001; Edwards et al., 2010). However, *CLCN2* is expressed in astrocytes and oligodendrocytes early in development (Makara et al., 2003) and has been detected in Bergman glia (Jeworutzki et al., 2012), which are important for neuronal migration in the formation of cortical structures. Together with our findings, these observations suggest that it would be worth exploring the role of this channel in the vertebrate neural stem cell niche. Interestingly, expression of *CLCN2* has been found outside the brain in an unrelated stem cell niche. It is expressed in Sertoli cells (Bösl et al., 2001), which are the primary somatic cells of the seminiferous epithelium that form the spermatogonial stem cell niche through physical support and expression of paracrine factors (Chen et al., 2005; Oatley et al., 2011). *CLCN2* mutant mice showed disorganized distribution of germ cells in tubules at 3 weeks, germ cells did not pass beyond meiosis I, and were completely lost at later stages (Bösl et al., 2001; Edwards et al., 2010). Hence, similarly to ClC-a regulation of proliferation in the neural stem cell niche, CLC-2 could be regulating proliferation in the spermatogonial stem cell niche.

Although the Sertoli *CLCN2* expression/germ cell depletion correlation in mouse is in accordance with our data suggesting an important role of the ClC-a/CLC-2 chloride channel in stem cell niches, it remains unclear how a chloride ion channel could non-autonomously modulate the mitotic activity of proliferative cells. ClC-a function in Malpighian tubules has been associated with the movement of Cl^−^ ions (Cabrero et al., 2014), but it is possible that its function in glia of the stem cell niche is unrelated to ion exchange. For example, it might recruit signaling molecules to modulate neuroblast proliferation. Conceptually, one way to test whether the channel function is related to the movement of ions would be to perform rescue experiments of *ClC-a* mutant phenotypes with a channel defective for the pore function. In practice, however, this type of experiment is not that straightforward since CLC-2 pore gating is quite complex. Channels of the CLC family are thought to function as a homodimers, with each subunit forming a pore and presenting both independent and common pore gating mechanisms (Jentsch and Pusch, 2018). Given the many studies supporting the function of CLC-2 as a channel, we next discuss different ways in which ionic imbalance caused by mutations in *ClC-a* could result in the phenotypes described. One of the possibilities we considered was whether ionic imbalance in *ClC-a* mutants affected secretion. Glial cells secrete different types of factor to the extracellular space, both during development and to maintain morphology in the adult (Coutinho-Budd et al., 2017; Read, 2018; Spéder and Brand, 2018). In the niche in particular, there are several examples of glia-secreted molecules that regulate neurogenic proliferation, such as the transforming growth factor a (TGF-a)-like ligand (Morante et al., 2013) and insulin-like peptides (dILPs) (Chell and Brand, 2010; Sousa-Nunes et al., 2011). In vertebrates, an increase in intracellular Ca^2+^ in astrocytes, which is caused by activation of G protein–coupled receptors and release of calcium from intracellular stores or calcium entry from the extracellular space through different types of channel, has been reported to evoke the release of gliotransmitters (Bazargani and Attwell, 2016; Khakh and McCarthy, 2015; Shigetomi et al., 2016). In this scenario, membrane potential changes mediated by Cl^−^ channel activity could modulate activation of GPCR or voltage dependent Ca^2+^ channels, mediating an increase in the Ca^2+^ intracellular concentration and resulting in secretion. In fact, the opening of voltage dependent Ca^2+^ channels has been proposed as the mechanism behind the increase in aldosterone production and secretion (Fernandes-Rosa et al., 2018) resulting from gain-of-function mutations of *CLCN2*, which are behind primary aldosteronism and cause sustained depolarization of glomerulosa adrenal cells (Fernandes-Rosa et al., 2018; Scholl et al., 2018). To test whether loss of function of ClC-a/CLC-2 channels also affected secretion, we performed glia-specific RNAi knock down of key upstream regulators of intracellular calcium release such as *Drosophila* IP3R and RyR receptors, and downstream effectors of calcium-regulated secretory vesicle exocytosis, as well as secretion assays in primary glial cultures where *CLCN2* was knocked down with RNAi (data not shown). However, we were unable to consistently recapitulate *ClC-a* mutant phenotypes or detect secretion defects, suggesting that if the absence or reduction of the channel impairs secretion, it does so only in a very limited way.

Another possibility is that ClC-a is involved in pH regulation. Under extracellular neutral pH, H^+^ and HCO_3_^−^ combine to form H_2_CO_3_, which in turn is in equilibrium with H_2_O and CO_2_. In acidic conditions, to compensate for the increase in H^+^, the HCO_3_^−^/Cl^−^ exchangers extrude HCO_3_^−^ to the extracellular space to form more H_2_CO_3_ and drive the reaction to the formation of H_2_O and CO_2_. Rat ClC-2 opens in response to extracellular acidification, allowing Cl^−^ to exit the cell (Jordt and Jentsch, 1997). Since for each molecule of HCO_3_^−^ extruded, one of Cl^−^ is internalized, ClC-2 activation might be required to regulate HCO_3_^−^ transport and allow the presence of extracellular Cl^−^, thus creating a Cl^−^ recycling pathway for HCO_3_^−^/Cl^−^ exchangers (Bösl et al., 2001). Assays in *Xenopus* oocytes have shown that ClC-a activity is also sensitive to pH (H. G-P. and R. E., unpublished results). Thus, it may be that the lack of ClC-a in cortex glia leads to a more acidic extracellular pH due to deficient Cl^−^ recycling for HCO_3_^−^/Cl^−^ exchangers. Since changes in extracellular and intracellular pH have been shown to affect the proliferative capacity of both wild type and cancer cells (Carswell and Papoutsakis, 2000; Ciapa and Philippe, 2013; Flinck et al., 2018; Persi et al., 2018; White et al., 2017), ClC-a function in pH regulation could explain the proliferation defects observed in the mutant.

Regardless of the molecular mechanism that mediates the effect of ClC-a on proliferation, our findings support the notion that glia-mediated ionic balance could be important for brain development. Our results are in accordance with those of recent studies suggesting a link between several ion channels and the development of the nervous system, with channels being important both in stem cells (Li, 2011; Liebau et al., 2013) and glia (Olsen et al., 2015). A recent example of a channel function in stem cells is the gene SCN3A, which codes for the NaV1.3 sodium channel. This channel is mainly expressed during development and is highly enriched in basal/outer radial glia progenitors and migrating newborn neurons (Smith et al., 2018). The appearance of this type of progenitor and defined neuronal migration has been associated with the establishment of gyri in the cortex (Fietz et al., 2010; Hansen et al., 2010; Reillo et al., 2011). Intriguingly, mutations in the SCN3A gene result in structural malformations of gyri in the cortex (Smith et al., 2018). Another example is the glial-specific Kir4.1 channel, which is related to neurodevelopmental disorders with associated cognitive defects. Mutations in *KCNJ10*, which codes for the glial-specific Kir4.1 channel, underlie SeSAME/EAST syndrome (seizures, sensorineural deafness, ataxia, intellectual disability and electrolyte imbalance/epilepsy, ataxia, sensorineural deafness and tubulopathy) (Bockenhauer et al., 2009; Scholl et al., 2009) and have been detected in patients diagnosed with autism spectrum disorder and epilepsy (Sicca et al., 2016, 2011). Reduced Kir4.1 expression in astrocytes significantly contributes to the etiology of Rett syndrome (Kahanovitch et al., 2018; Lioy et al., 2011), which shares many pathophysiological traits with SeSAME/EAST. Moreover, Kir4.1 protein is detected as early as embryonic day 20 in glial cells in the developing cortex and hippocampus (Moroni et al., 2015), suggesting that it could influence neural development in these regions. Together with our findings, these observations suggest that mutations in ion channels could affect neurogenesis and connectivity, resulting in intellectual disabilities. Thus, providing insights into the developmental stages affected by impaired glial-dependent homeostasis could help improve our understanding of the origin of the cognitive deficiencies detected in patients with channelopathies or conditions where ion channels in glia are not functional.

## MATERIALS AND METHODS

### Genetics

Flies were grown in standard medium at 25°C except for RNAi experiments, which were performed at 29°C. All genotypes analyzed are specified in the Supplementary Information.

Stocks used to characterize *ClC-a* expression and phenotype were: *MiMIC 05423* (Bloomington Drosophila Stock Center, BDSC 43680), *05423^ClC-a-GAL4^* (BDSC 66801), *MiMIC 14007* (BDSC 59247), *Df(3R)PS2* (BDSC 37742), *mir8-GAL4* (DGRC 104917), *R54H02-GAL4* (BDSC 45784), *wrapper932i-LexA*, *wrapper932i-GAL80* (Coutinho-Budd et al., 2017), *repo-GAL4* on II (Lee and Jones, 2005), *repo-GAL4* on III (BDSC 7415), UAS-Dcr2 (Vienna Drosophila Resource Center, VDRC 60009), *UAS-ClC-a-RNAi* (VDRC 110394), *UAS-ClC-a* and *UAS-ClCN2* (this study), *UAS-slit-RNAi* (VDRC 108853), *slit^dui^* (BDSC 9284), *Slit-GFP* (BDSC 64472), and R9D11-tdtom (BDSC 35847). Additional stocks used in Supplementary Figures were: *ClC-a-GFP* (BDSC 59296), *slit-lacZ* (*Slit^05428^)* (BDSC 12189), *Rh1GAL4* (BDSC 68385), *Rh4EGFP* (BDSC 7462), *Rh6-lacZ* (BDSC 8117), *GMR-GAL4* (BDSC 1104), *R43H01-LexA* (BDSC 47931) and *R25A01-GAL4* (BDSC 49102), *gcm-lacZ* (BDSC 5445).

To label membranes and nucleus, we used *UAS-mCD8-GFP* (BDSC 5137), *UAS-mCD8-RFP.LG* (BDSC 27398), *UAS-mCD8GFP,lexAop-CD2RFP* (BDSC 67093), *UAS-H2B-RFP* (Mayer et al., 2005), and *UAS-H2B-YFP* (Bellaïche et al., 2001), as specified in the genotype list. In experiments where nuclear labeling was used for quantification, the same transgene was employed for control and mutant samples (Fig 5M, Sup Fig 5A).

To generate and label neurogenic tissue clones in control and *ClC-a* mutant backgrounds (Figure 4D,E and Figure 8J), the following stocks were crossed: *hsFLP,FRT19A,tub-Gal80; tub-GAL4,UAS-mCD8GFP/CyODfYFP; 14007/+* to *FRT19A; +; +* and *hsFLP,FRT19A, tub-Gal80; tub-GAL4,UAS-mCD8GFP/CyODfYFP; 14007/+* to *FRT19A; +; Df(3R)PS2/TM6B*. Three-hour egg lays were maintained at 25°C and clone induction was performed with a 30-minute heat shock pulse at 37°C in a water bath at the L2 stage. Brains were dissected 48 hours after clone induction.

For lineage-tracing experiments (Supplementary Figure 8D-F, G), we used G-TRACE (*UAS-RedStinger,UAS-FLP,Ubi-FRT-stop-FRT-Stinger*, BDSC 28280 (Evans et al., 2009) combined with specific *GAL4* drivers.

When we performed the cortex glia-specific rescue experiments (Figure 4J, K and Figure 7B, C), no cortex glia-specific driver had yet been published, so we devised an intersectional genetic strategy to generate one using the *mir8-GAL4* driver. In addition to surface associated cortex glia on the OPC, *mir8-GAL4,* labels cortex glia and neurons in the brain, as well as other cells in the animal. To restrict *mir-8* expression exclusively to cortex glia, we combined the following transgenes: *repo-FLP6.2* (Stork et al., 2014), *tub>GAL80>* (BDSC 38879), and *mir8-GAL4*. In this combination, GAL4 is only expressed in cortex glia since the GAL80 repressor has only been flipped out in this cell type but persists in non-glial mir-8 expressing cells (Supplementary Figure 7). For the sake of simplicity, we refer to this combination as the *mir-8 ^cxg^* driver.

### DNA constructs

For *UAS-ClC-a* and *UAS-CLCN2* transgenes, we used the Gateway cloning system (Invitrogene) and cloned their respective cDNAs, to which a 3xFLAG tag had been previously added in the C-terminus, into the ΦC31 integrase compatible pBID-UASC-G plasmid (Addgene plasmid # 35202, a gift from Brian McCabe (Wang et al., 2012)). The FLAG tag does not alter the electrophysiological properties of these channels. For the *ClC-a* construct, we used the isoform C (a gift from P. Cid) since its electrophysiological properties have already been studied in *Xenopus* oocytes and HEK-293 cells (Flores et al., 2009; Jeworutzki et al., 2012), and it is known to be expressed in *Drosophila* head and body (Flores et al., 2009). The final constructs were injected into the *attp40* (25C6) landing site on the 2^nd^ chromosome.

### Immunohistochemistry

Fly brains were dissected in Schneider medium and fixed in 4% PFA in PBL (75 mM lysine, 37 mM sodium phosphate buffer, pH 7.4) for 25 min. After fixation, the tissue was washed in PBS with 0.5%Triton-X-100 (PBST) and blocked with PBST with 10% normal goat serum. Primary and secondary antibody incubations were performed in PBST and 10% normal goat serum, typically overnight at 4°C. The following primary antibodies were used for immunohistochemistry: mouse anti-Chaoptin (1:50, 24B10, Developmental Studies Hybridoma Bank, DSHB), rat anti-DE-cadherin (1:50, DCAD2, DSHB), mouse anti-Repo (1:50, 8D12, DSHB), rabbit anti-ClC-a (1:100 this study, see Supplementary Information, and 1:100 a gift from J. Dow), guinea pig anti-Deadpan (1:2000, gift from A. Carmena), rat anti-Lethal of Scute (1:5000, gift from A. Brand), rabbit anti-Mira (1:500, gift from C. González), chicken anti-GFP (1:800, ab13970, Abcam, Cambridge, UK), rabbit anti-RFP (1:200, 632496, Clontech, Mountain View, CA, USA), mouse anti-β-galactosidase (1:1000, Z3783, Promega, Madison, WI, USA), rabbit anti-β-galactosidase (1:1000, 0855976, Cappel, Malvern, PA, USA), and rabbit anti-Dcp-1 (1:200, Asp216, Cell Signaling Technology, Danvers, MA, USA). Alexa Fluor 488, 568, and 647 secondary antibodies raised in rabbit, mouse, rat, guinea pig, or chicken (Life Technologies, Carlsbad, CA, USA) were used at 1:250 concentration. Nuclei were labeled using TOPRO-3 (1:1000, T3605, Life Technologies). Brains were mounted for confocal microscopy in Vectashield (Vector Laboratories, Burlingame, CA, USA). Samples were analyzed with Leica TCS SPE and Zeiss LSM880 confocal microscopes.

### Photoreceptor phenotype classification

We classified brains as having a strong, medium, or weak photoreceptor phenotype based on the OL that out of the two had the most severe phenotype. If not the same, the two OLs tended to be in consecutive categories (i.e. strong/medium, medium/weak, weak/no phenotype). For experiments involving photoreceptor phenotypes, we always analyzed at least 17 brains.

### Measurements and quantifications

To assess adult OL and CB size, we took two confocal images of each brain in the appropriate orientations to measure the antero-posterior and dorso-ventral axis of each structure at their widest part. We multiplied those two measurements to obtain a relative value in arbitrary units (a.u). The number of brains analyzed ranged from 23 to 44 for OL (to obtain fully independent measurements, only one OL per brain was quantified), and from 12 to 22 for CB when assessing phenotype (Figure 2C, Figure 4H, I). In rescue experiments (Figure 2D, Figure 4J, K), the number of brains analyzed ranged from 12 to 32.

To assess brain size at different larval stages (Figure 4G), the diameter of one larval brain hemisphere per animal was measured in the antero-posterior axis. The *n* ranged from 23 to 37 animals analyzed.

To quantify the number of cortex glia nuclei in late L3 OLs, we manually counted *ClC-a*^+^ nuclei (Supplementary Figure 5A). Cortex glia nuclei present an average size of approximately 44.5 μm^2^ (Morante et al., 2013) and are clearly distinguishable from ClC-a^+^ ensheathing glia nuclei by their size and position in the OL. The *n* for this experiment was 8 brains.

To count the number of neuroblasts in the CB part of the hemisphere (Supplementary Figure 5B), we manually identified them based on their nuclear size and on Dpn and Mira antibody staining. Neuroblasts were distinguishable from mature INPs, which are also Dpn^+^ and Mira^+^, by their smaller nuclear size and higher intensity of TOPRO-3 nuclear staining. The *n* for this experiment was 8 brains. To measure neuroepithelia volume, the tissue was stained with anti-E-cadherin antibody and manually segmented using the “SURFACE” tool included in Imaris software. This tool provides the volume in μm^3^ of the surfaces generated (Figure 4B). The *n* for this experiment was between 8 and 9 brains.

To quantify the number of cells in OPC, IPC, and CB neuroblast clones (Figure 4D, E), we performed manual counting in confocal stacks. Cells in the clone were identified as TOPRO+ nuclei surrounded by labeled membrane. We counted as many clones as possible per brain provided that they were identifiable as individual clones. The number of clones analyzed was 52 (control) and 31(mutant) in the OPC and 18 (control and mutant) in the IPC, and the number of type I neuroblast clones was 39 (control) and 22 (mutant).

To assess cell death in developing OLs (Figure 4F), we manually counted Dcp-1^+^/TOPRO + puncta per brain hemisphere. This value was divided by the hemisphere volume obtained through manual segmentation of the structure and using the “SURFACE” tool included in Imaris software. The *n* for this experiment was 6 brains. To quantify the subset of boundary glia among glial cells in the chiasm region at different stages, we manually counted ClC-a^+^/Repo^+^ nuclei (Figure 5M). The *n* for this experiment was 5 brains.

To quantify mature INPs in the DL1 and DL2 lineages (Figure 5U), we manually counted Dpn^+^ positive nuclei surrounded by tdtom^+^ membranes of the *R9D11-tdtomato* marker. To differentiate DL1 from DL2, we used *gcm-lacZ*, which specifically labels the DL2 lineage (Supplementary Figure 9). The *n* for this experiment was between 11 and 12 brains.

To quantify the number of total glial cells in the future chiasm region in mid L3 (Figure 7B), we manually counted Repo^+^ nuclei in confocal stacks. The *n* for this experiment was between 5 and 8 brains.

### Image processing

Fiji or Imaris 8.0 (Bitplane, South Windsor, CT, USA) were used to process confocal data as specified in figure legends. Figures were assembled using Adobe Illustrator (Adobe, San Jose, CA, USA).

### Statistics

Statistical analysis was carried out using Prism 6 (GraphPad Software Inc, San Diego, CA, USA). When data did not follow a normal distribution or resulted from a previous mathematical computation (i.e. ratio to volume), we used non-parametric tests. For group comparisons, we used Kruskal-Wallis followed by planned pairwise comparisons with the Mann-Whitney post hoc test to obtain *p*-values (Figure 2C, D; Figure 4B, H, I; Figure 5M; Figure 7B; Supplementary Figure 8H). For pairwise comparisons, we used Mann-Whitney’s test (Figure 4D, E, F, J, K, Figure 5U, Supplementary Figure 9B). For paired comparisons, we used the Wilcoxon matched-pairs squad rank test (Figure 5U). Data are shown in box plots where the median is given between the first and third quartiles. Whiskers represent the maximum and minimum values of the data. When all data in the analysis were suitable for a parametric analysis, we performed one-way ANOVA followed by Turkey’s post hoc test to obtain *p*-values (Figure 4G). *p*-values for pairwise comparisons relevant to our biological inquiry are shown in the bar graph. Data is represented as the mean, and error bars show standard deviation.

To evaluate the statistical significance of enhancements in qualitatively categorized photoreceptor phenotypes (i.e., strong, medium, weak, no phenotype) (Figure 6L, M), we performed a Chi-squared test of independence between phenotype categories and genotypes to obtain *p*-values.

## ACKNOWLEDGMENTS

We are grateful to A. Brand, H. Bellen, A. Carmena, P. Cid, J. Coutinho-Budd, J.A. Dow, M. Freeman, C. Gonzalez, B.W. Jones, C. Klambt, J. Morante, I. Salecker, M. Wernet, DSHB, BDSC, and VDRC for reagents. We thank V. Hartenstein, J. Modolell, C. González, A. Carmena, J. Tejedor, and C. Homem for helpful discussions and suggestions on optic lobe and brain development. We thank F. Aguado, N. Barranco, and X. Elorza-Vidal for performing secretion experiments in cell culture. We thank M. Bosch from the Confocal Unit of the CCiT-UB. We thank F. Cebriá for critical reading of the manuscript. Our gratitude to M. Corominas, F. Serras, and members of their laboratories for engaging in discussions and making suggestions during our joint lab meetings throughout the project. This work was funded in part by the Spanish Ministry of Economy and Competitiveness through grants BFU2015-69689-P to M.M. and M.R, and SAF2015-70377 to R.E.; the Generalitat de Catalunya through grant SGR2014-1178 to R.E.; the Institució Catalana de Recerca i Estudis Avançats through an ICREA Academia award to R.E.; and the University of Barcelona through the award of an APIF fellowship to H.P-S.

### AUTHOR CONTRIBUTIONS

M.M. and R.E. conceived the project; M.M., H.P-S., Q.Z., R.E., and H. G-P. designed the experiments and data analysis; H. G-P. and R.E. contributed reagents and analytical tools; M. R. designed the statistical analysis; H.P-S. and Q.Z. performed the experiments; H.P-S., Q.Z, M.R., and M.M. analyzed the results; and M.M. wrote the manuscript with contributions from all the other authors.

