## Supplementary for "*Drosophila ClC-a* is required in glia of the stem cell niche for proper neurogenic proliferation and wiring of neural circuits"

### SUPPLEMENTARY INFORMATION

This file contains
-Supplementary Material and Methods
-Supplementary Figures and Legends
-References
-Genotype list

### Supplementary Material and Methods

#### Genetics

ClC-a-GFP: As with *ClC-a-GAL4*, the ClC-a-GFP line we used was also derived from *Mi(MIC)ClC-a*$^{05423}$ by the Gene Disruption Project. The protein trap cassette used to generate ClC-a-GFP contains the GFSTF tag (EGFP-FlAsH-StrepII-TEV-3xFlag) flanked by flexible linkers on both sides in the appropriate phase. The ClC-a tagged protein generated includes the GFP sequence in frame in an extracellular loop of the protein. Homozygous animals are viable and wild type, indicating that the ClC-a-GFP protein is functional.
*ClC-a* alleles: Viability analysis of MiMIC insertions over deficiency revealed that the very few *05423/Df* animals that emerged from the pupal case did so 48 hours later than heterozygote controls, and remained immobile on the food before dying shortly after eclosure. Escapers of the *14007/Df* allelic combination emerged with a delay of around 24 hours, but compared to *05423/Df* were more abundant and healthier, all emerging from the pupal case. Similar to *14007/Df* animals, *05423*$^{ClC\text{-}a\text{-}GAL4}$*/14007* also had a 24-hour developmental delay with respect to controls. This developmental delay has also been confirmed with a morphological developmental landmark: photoreceptor innervation. The third instar larval stage (L3) can be divided into early, mid, and late stages. In wild type and heterozygous control animals, photoreceptor innervation takes place at mid L3 (96 hours after egg lay (AEL)) and pupariation at late L3 (120 hrs AEL). However, in *14007/Df* and *05423*$^{ClC\text{-}a\text{-}GAL4}$*/14007* mutant animals, photoreceptors did not enter the brain until 120 hrs AEL and pupariation did not take place until 144 hrs AEL. Glia-specific rescue experiments also rescued the developmental delay. To compare control and mutant animals at the same developmental stage, we took developmental delay into account, and for the sake of simplicity, we refer to comparisons in larval stages as opposed to developmental hours (i.e., we refer to comparisons as control versus mutant at mid L3 rather than 96 hrs AEL control versus 120 hrs AEL mutant). In all cases, adult mutant animals were of the same size as heterozygote controls and wild type flies (data not shown).
Slit-GPF: similar to ClC-a-GFP, this Slit protein trap was derived from *Mi(MIC)sli*$^{03825}$ by the Gene Disruption Project.

#### Antibody generation

Immune sera against synthetic peptides from *Drosophila melanogaster* ClC-a (RVIDMSPEDQKQWEL, corresponding to amino acids 874-888, and ESKQSPSADKSNTENGNHA, corresponding to the last 19 amino acids of the protein, 1031-1049) were raised in rabbits using the services provided by Eurogentec. Peptides were coupled to keyhole limpet hemocyanin via a cysteine residue that had been added to the C-terminal of the peptide. After four/five boosts of immunization, the antisera were affinity purified using the peptide covalently coupled to Sulpholink (ThermoScientific). The polyclonal antibody was tested by immunoblotting in HEK293A cells transfected with the pcDNA3.1 vector (Invitrogen) expressing the ClC-a channel with a 3xFLAG-tag fused to the

C-terminus.

**Western blots**

Two different protein extraction procedures were used. For the antibody testing, HEK293A cells were grown on DMEM containing 10% (v/v) fetal bovine serum (Sigma) and 1% penicillin/streptomycin at 37ºC in a humidity controlled incubator with 10% $CO_2$. Cells were transfected with 2 µg of the *ClC-a 3xFLAG* pcDNA3.1 construct using the Transfectin reagent (BioRad). Forty-eight hours after transfection, cells were harvested and homogenized in lysis buffer containing 1% TX-100, 150 mM NaCl, and phosphate-buffered saline plus protease inhibitors as described elsewhere (Capdevila-Nortes et al., 2013). In the case of fly tissue protein extracts, twenty *Drosophila* heads were homogenized by 20 strokes in an Eppendorf Teflonglass homogenizer in 10 µl/brain RIPA buffer containing 50 mM TRIS/HCl pH8, 150 mM NaCl, 1% NP40, 0.5% deoxycholate, 0.1% SDS, 2 mM EDTA, and protease inhibitors (aprotinin (1 µg/ml), PMSF (174.2 µg/ml), Leupeptin (1 µg/ml), and Pepstatin (1 µg/ml)). For both extracts, the proteins in the supernatant were quantified by the BCA method. Sodium dodecyl sulfate polyacrylamide gel electrophoresis and western blot were performed as described elsewhere (Teijido et al., 2004) loading 50 µg of *Drosophila* head extracts, or 100 µg of HEK293A cell extracts. We used our custom polyclonal antibody to detect ClC-a (1:100 dilution), mouse monoclonal anti-Flag M2 (Sigma-Aldrich) (1:500 dilution), and anti-Tubulin (MMS-435P, Covance Antibody Products, Princeton, NJ) (1:500 dilution). Secondary antibodies were horseradish peroxidase-conjugated anti-rabbit and anti-mouse (Jackson ImmunoResearch).

### Supplementary Figures and legends

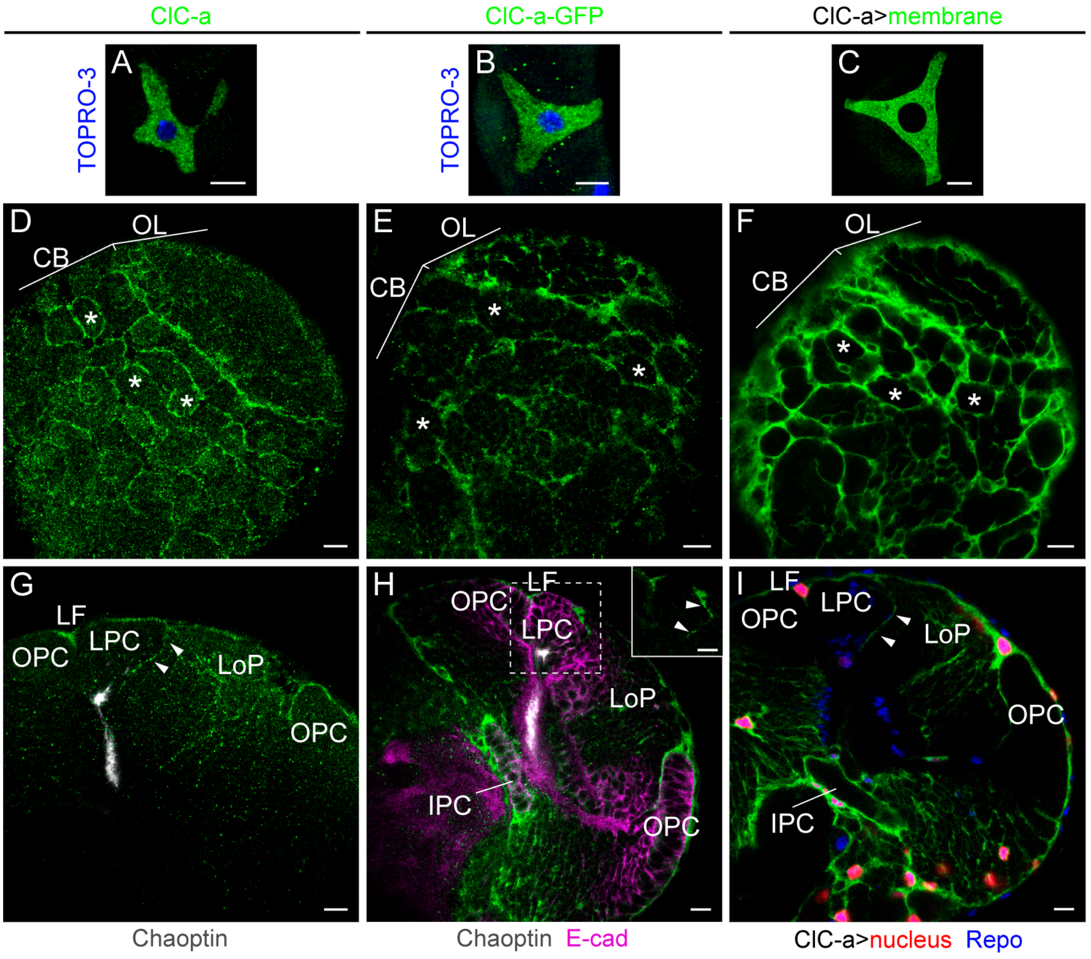


**Supplementary Figure 1. Comparative analysis of *ClC-a* expression patterns with antibody and various reporters.**

(A-C) Detection of *ClC-a* expression (green) in stellate cells of adult Malpighian tubules using anti-ClC-a antibody (A), the *ClC-a-GFP* protein trap (B), and the *ClC-a-GAL4* driver line combined with a membrane reporter (green) (C). Nuclei labeled with TOPRO-3 where indicated. (D-I) Detection of *ClC-a* expression in late L3 brain hemispheres. (D-F) Horizontal views of the surface of brain hemispheres. Antibody staining (D), protein trap (E), and driver (F) show the same expression patterns. Asterisks mark some neuroblast chambers. (G-I) Horizontal views deeper in hemispheres, in the optic lobe area. Arrowheads point to ClC-a expression between the LPC and LoP. (G) Antibody staining shows expression on the OPC, in the LF, and in between the LPC and LoP. (H) In addition, the protein trap construct also reveals expression deeper in the brain, around the IPC and forming a mesh-like structure inside the hemisphere, where the antibody did not penetrate. Inset shows expression between the LPC and LoP. Anti-E-cad staining (magenta) was used to identify the neuroepithelial cells and anti-Chaoptin (24B10, gray) label photoreceptors. (I) The *ClC-a-GAL4* driver mediated membrane labeling (green) pattern is very similar to the one observed with the antibody and the protein trap construct, including the signal detected between the LPC and LoP. Glial nuclei were labeled with anti-Repo antibody (blue). Not all glial nuclei are ClC-a$^{+}$ (red).

CB, central brain; OL, optic lobe; LF, lamina furrow; LPC, lamina precursor cells; LoP, lobula plug; OPC, outer proliferation center; IPC, inner proliferation center. Scale bars represent 10 µm.

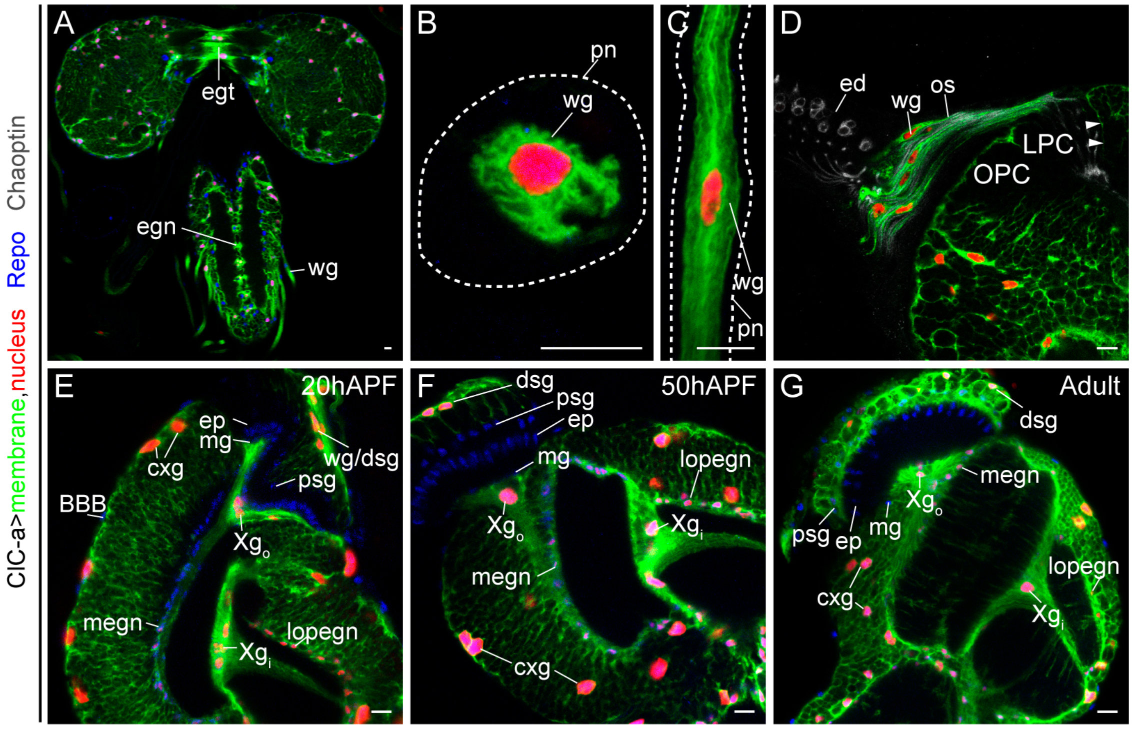


**Supplementary Figure 2. Identification of *ClC-a* expressing glia.**

Confocal sections showing *ClC-a* expression pattern in the late L3 nervous system (A-C), and the optic lobe in pupal stages (D, E) and adult (F). *ClC-a* specific GAL4 driver was used to label cellular membranes (green) and nuclei (red) of ClC-a$^{+}$ cells. Glial nuclei were labeled with anti-Repo antibody (blue) and photoreceptor cells with anti-Chaoptin (gray). (A) Larval brain where, besides a ClC-a$^{+}$ signal in cortex glia both in brain hemispheres and the VNC, a ClC-a$^{+}$ signal is detected in neuropil-ensheathing glia in the VNC, tract-ensheathing glia in connectives between the two hemispheres, and in peripheral nerves. (B,C) Cross section (B) and longitudinal section (C) of peripheral nerves containing ClC-a$^{+}$ glia. Dashed line outlines the nerve. (D) Image of the optic stalk, which connects the eye disc and the optic lobe. ClC-a$^{+}$ glia wraps this bundle formed by photoreceptor axons on their way to the optic lobe. Photoreceptor cell bodies are seen in the eye disc in gray and their axons in the optic lobe. Photoreceptors do not express *ClC-a*. (E, F) Based on the ClC-a$^{+}$ Repo$^{+}$ nucleus position, we can identify the following as *ClC-a* expressing glia: cxg, wg/dsg, $Xg_o$, $Xg_i$, mneg, and lopneg. (G) *ClC-a* expression is maintained in the adult. Signal in the medulla and lobula neuropils belongs to mneg and lopneg described projections into these structures.

egt, tract-ensheathing glia; egn, neuropil-ensheathing glia; wg, wrapping glia; pn, peripheral nerve; ed, eye disc; os, optic stalk; OPC, outer proliferation center; LPC, lamina precursor cells; BBB, blood brain barrier; ; cxg, cortex glia; megn, medulla neuropil-ensheathing glia; ep, epithelial glia; mg, marginal glia; $Xg_o$, outer chiasm glia; $Xg_i$, inner chiasm glia; psg, proximal satellite glia; wg/dsg, wrapping glia/distal satellite glia; lopegn, lobula plate neuropil-ensheathing glia. Scale bars represent 10 µm.

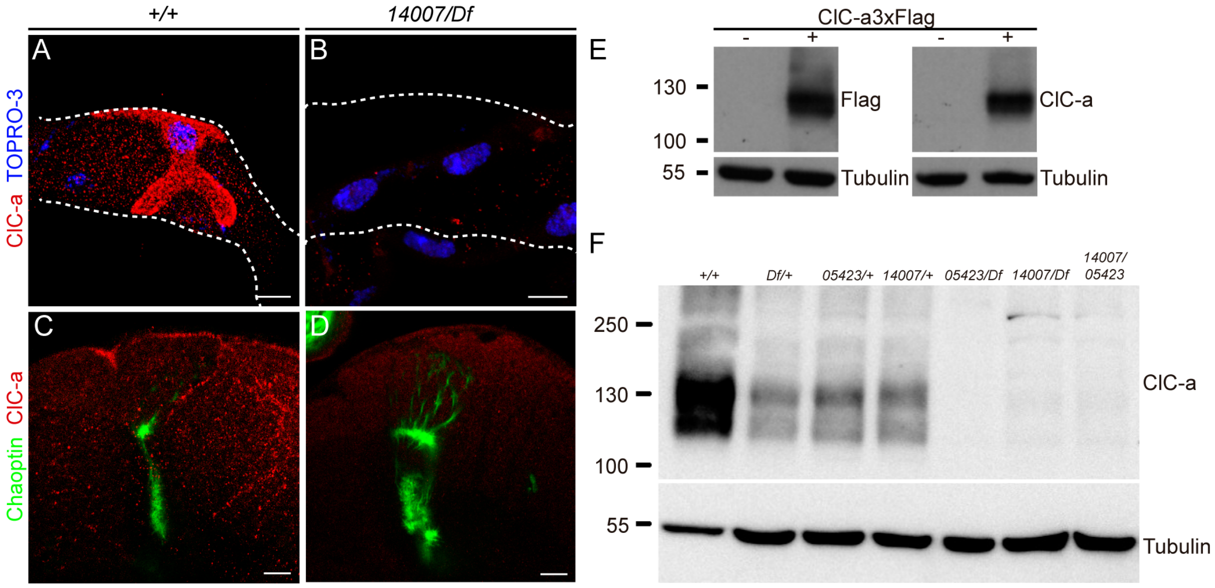


**Supplementary Figure 3. Immunohistochemistry and western blot analysis of *ClC-a* MiMIC alleles.**

(A, B) Anti-ClC-a antibody staining of adult Malpighian tubules in control animals (A) and *14007/Df* mutants (B). (C, D) Anti-ClC-a antibody staining of late L3 brains in control animals (C) and *14007/Df* mutants (D). Photoreceptors are labeled with anti-Chaoptin (24B10, green). (E) Western blot of protein extraction from HEK293 cells transfected with or without *ClC-a isoform C 3xFlag* pcDNA3.1. Both anti-Flag and anti-ClC-a antibodies detect a band below 130 kDa, which is possibly the weight of the protein (Uniprot prediction at 118 kDa) plus glycosylation. (F) Western blot of protein extraction from adult heads of controls and different allelic combinations. The signal around the 130 kDa mark reflects the presence of ClC-a protein in controls, most probably of different isoforms which range from 113 to 132 predicted kDa plus glycosylation. A strong reduction in this signal is observed in mutant animals.

Scale bars represent 10 µm.

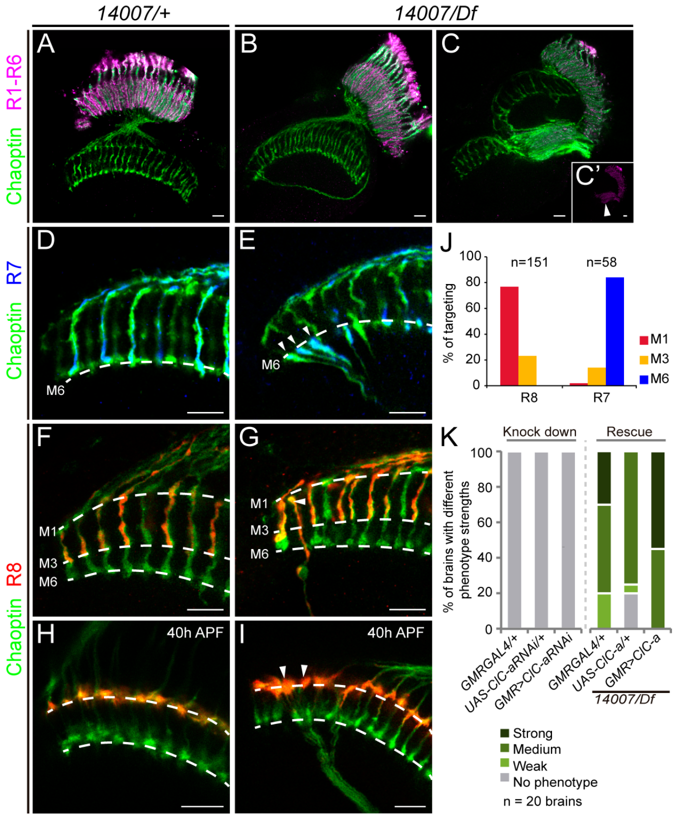


**Supplementary Figure 4. Analysis of layer selection defects in misguided photoreceptors of *ClC-a* mutants.**

Confocal images of adult (A-G) and pupal (H,I) optic lobes stained with anti-Chaoptin to label all photoreceptors (green). Photoreceptor subtypes were labeled using cell type specific opsin reporters: R1-6 (magenta), R7 (blue), and R8 (red). *An R8 specific driver (senseless)* was used to label R8s in pupal brains (red). (A) Control photoreceptor array showing R1-6 photoreceptors stopping in the lamina. (B, C) Mutant arrays. (B) In mutant animals with weak guidance defects, R1-6 terminate normally in the lamina. (C) In animals with strong guidance defects, R1-R6 axons invade the medulla as seen in the inset (C'). (D) Control array showing R7s terminating at the M6 layer. (E) Mutant array shows misguided R7s terminating in the M6 layer like controls. (F) Control array showing R8s terminating in the M3 layer. (G) Mutant array showing misguided R8s terminating in the M1 layer. (H) Control array at 40 hrs pupal development. R8 cells terminate in the prospective M1 layer at the top of the medulla. This is a temporary stop since in a second stage they actively extend to the M3 layer. (I) Misguided R8s in the mutant animal also terminate in the M1 layer; however, the adult phenotype suggests that these cells are unable to detach from this temporary layer and retract to the M3. (J) Quantification of adult targeting defects in misguided photoreceptors. Most R8s terminate in M1 (red) instead of at M3, while most R7s terminate correctly at M6. The limited number of R7 targeting defects can be explained by the fact that in pupal stages, R7s already extend to a deeper layer with their growth cones very close to their synaptic partners, and that the R7 axons grow by intercalation of ingrowing processes of other neurons. (K) Quantification of the percentage of brains with different strengths of guidance phenotypes in photoreceptor-specific knock down and rescue experiments. Consistent with the absence of *ClC-a* expression in photoreceptors, misguidance phenotypes are non-autonomous. Eye specific *ClC-a* knockdown results in proper photoreceptor guidance and eye specific *ClC-a* expression in mutants does not rescue photoreceptor guidance phenotypes.

Scale bars represent 10 µm.

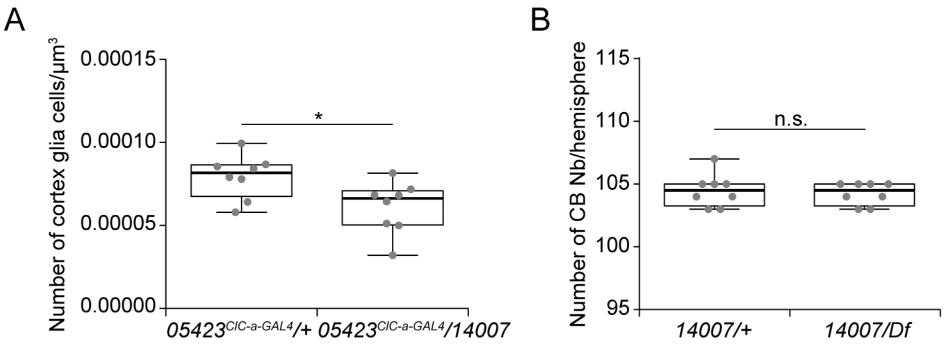


**Supplementary Figure 5. Quantification of ClC-a$^{+}$ cortex glia nuclei and central brain neuroblasts in control and *ClC-a* mutant brain hemispheres.**

(A) Ratio of cortex glia nuclei/µm$^3$ in late L3 control (*05423$^{ClC\text{-}a\text{-}GAL4}$*/+) and mutant (*05423$^{ClC\text{-}a\text{-}GAL4}$/14007*) brain hemispheres. (B) Quantification of the number of CB neuroblasts present in late L3 control (*14007/+*) and mutant (*14007/Df*) brain hemispheres. *p*-value was calculated with the non-parametric Mann-Whitney test. n.s.>0.05, *p<0.05.

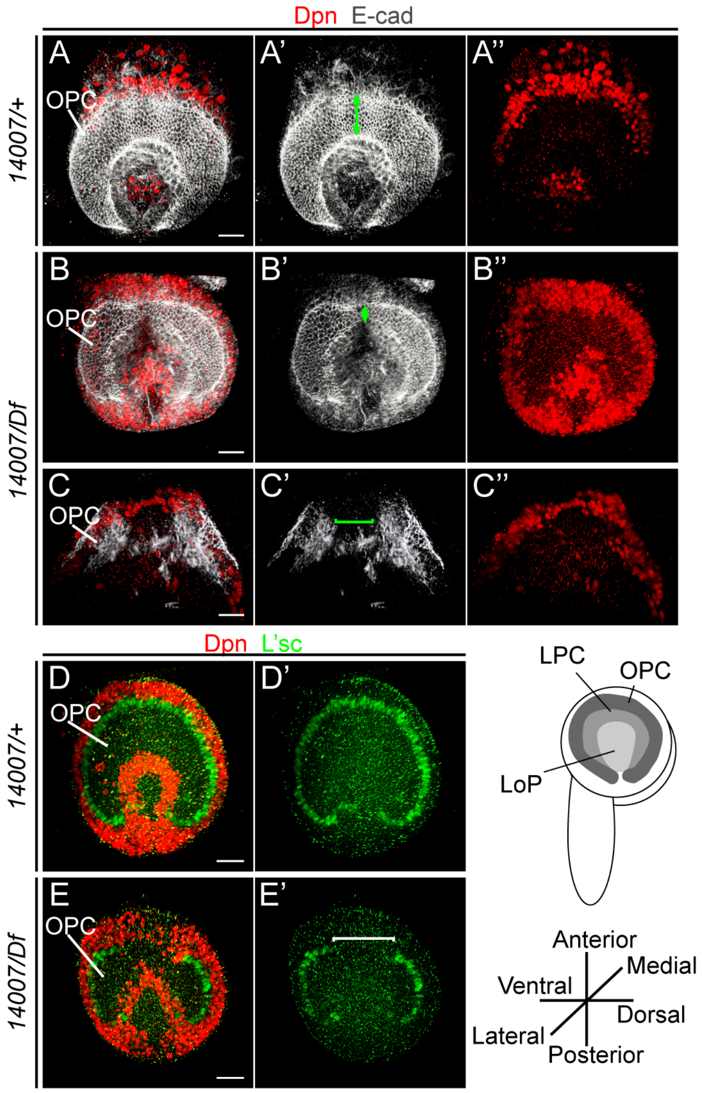


**Supplementary Figure 6. Analysis of neuroepithelium to neuroblast transition in *ClC-a* mutant animals.**

Lateral views of volume-rendering 3D reconstructions of late L3 larval hemispheres. (A) Control animal (*14007/+*) stained with anti-E-cad (gray, A') labeling the OPC and anti-Dpn (red, A'') labeling neuroblasts, which are differentiating on the medial side of the OPC. Double arrow marks the width of the OPC in the central region. (B, C) Examples of OPC defects observed in late L3 mutant hemispheres (*14007/Df*). (B) E-cad staining (B') reveals a reduction in the width of the OPC (double arrow), especially in the central part. (C) In this severe example, although E-cad staining is gone (bracket), there are neuroblasts, suggesting that the neuroepithelial to neuroblast transition in this region of the OPC took place prematurely and there is no more OPC tissue. (D) Control animal (*14007/+*) stained with anti-L'sc (green), which labels the neuroepithelial cell that will transition to neuroblast, and anti-Dpn (red) to visualize neuroblasts. (E) Mutant animal that lacked L'sc expression in the central region of the OPC. The absence of L'sc indicates that there was no more neuroepithelium to differentiate into neuroblasts. The presence of neuroblasts (red) in the region where L'sc is missing indicates that there used to be neuroepithelium.

Scale bars represent 10 µm.

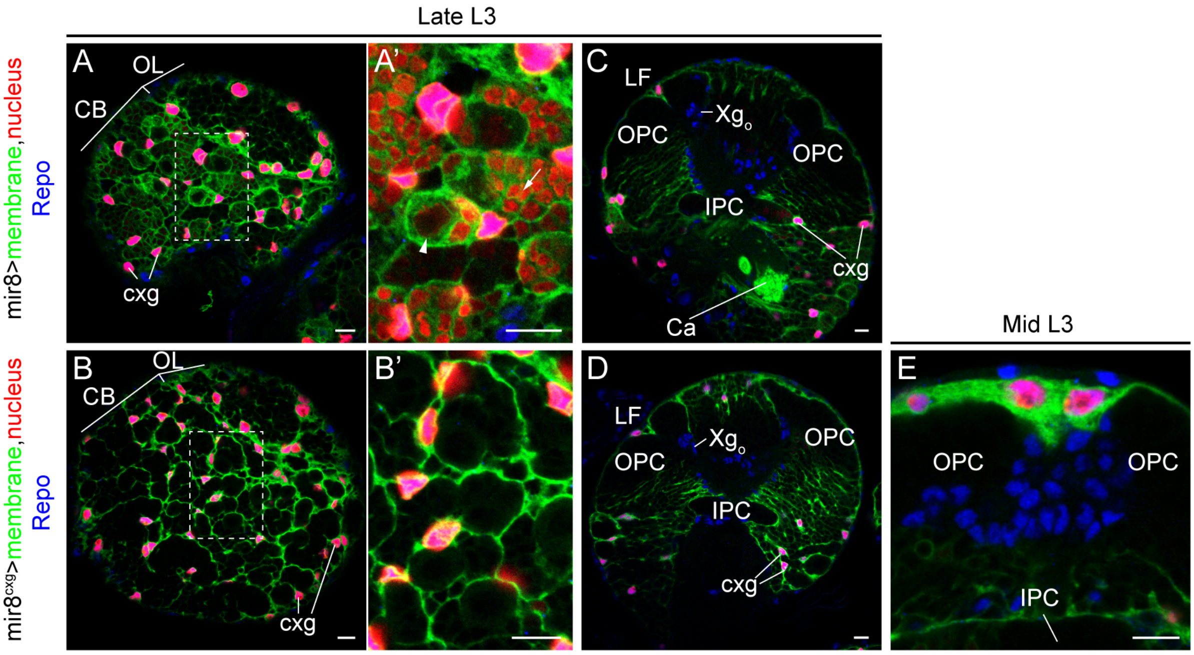


**Supplementary Figure 7. Characterization of a cortex glia-specific driver.**
Expression patterns of *mir-8-GAL4* driver and *mir-8* $^{cxg}$. Membranes were labeled in green, nuclei in red, and all glial nuclei in blue (anti-Repo). (A, B) Horizontal views at the surface of the central brain showing *mir-8* (A) and *mir-8* $^{cxg}$ expression (B). (A', B') Magnifications of dashed region of interest in (A) and (B). (A') *mir-8-GAL4* is expressed in neuroblasts (arrowhead) and neurons (arrow). The gain of the red channel has been increased to visualize nuclear signal in neuroblasts and neurons. (B') Magnification of dashed region of interest in (B). Using the same gain as in (A'), neuronal and neuroblast labeling is gone using the *mir-8* $^{cxg}$ transgenes. (C, D) Horizontal views deep in the brain hemisphere showing *mir-8-GAL4* (C) and *mir-8* $^{cxg}$ expression (D). (C) Neuronal *mir-8* expression is seen in the mushroom body calyx. $Xg_o$ glia do not express *mir-8*. (D) No neuronal expression was detected in the calyx or $Xg_o$. (E) Frontal view of a volume-rendering 3D reconstruction of a mid L3 optic lobe. No membrane (green) and/or nuclear (red) signal between the OPC and IPC confirmed that *mir-8* $^{cxg}$ was not expressed in boundary glia.
CB, central brain; OL, optic lobe; cxg, cortex glia; LF, lamina furrow; OPC, outer proliferation center; IPC, inner proliferation center; $Xg_o$, outer chiasm glia; Ca, calyx. Scale bars represent 10 μm.

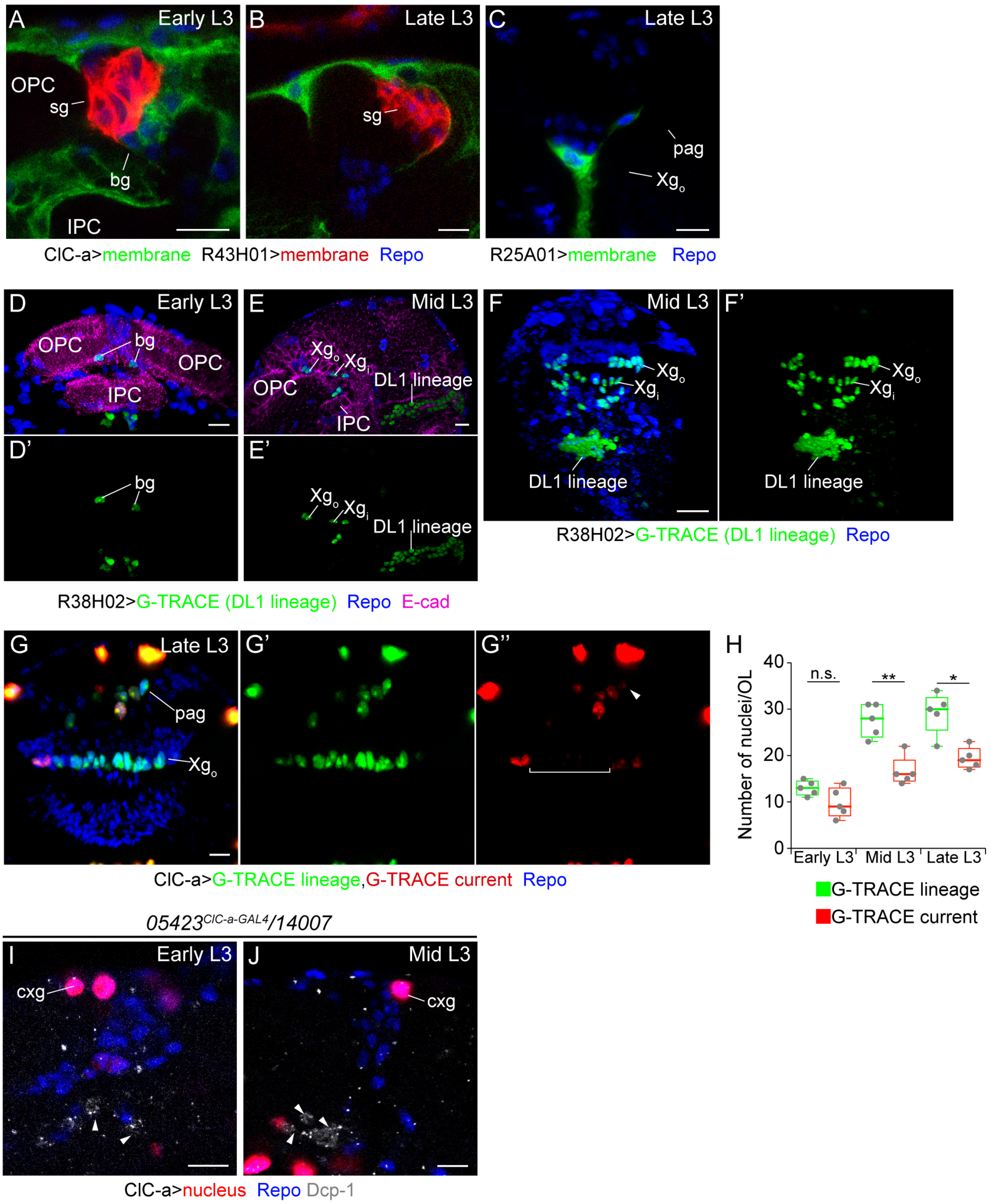


**Supplementary Figure 8. Developmental details of the formation of the glial barrier between the LPC and the LoP.**

(A-C) Characterization of cell types in the barrier. Specific drivers were used to label membranes in green or red. Glial nuclei are labeled with anti-Repo (blue). (A-B) Horizontal views of early (A) and late (B) optic lobes showing ClC-a$^-$ satellite glia population membranes labeled with the *R43H01-LexA* specific driver in red and ClC-a$^+$ membranes (*05423$^{ClC\text{-}a\text{-}GAL4}$* /+) in green. (C) Horizontal view of a late L3 optic lobe showing $Xg_o$ and palisade glia membranes labeled with the specific driver *R25A01-GAL4* in green. This driver is not expressed at earlier developmental time points, and thus cannot be used to manipulate these cell types when they group together as boundary glia before photoreceptor innervation in mid L3. (D-F) DL1 lineage tracing to analyze parallelisms between the timing of visualization of DL1 derived $Xg_o$ glia and visualization of ClC-a$^+$ boundary glia (prospective

$Xg_o$ and pag) in the optic lobe. DL1 lineage (green) is visualized with the DL1 specific driver *R38H02-GAL4*, which is expressed in this NB early in development in a short time window, and the G-TRACE system. Optic lobes were stained with anti-E-cad (magenta) to identify neuroepithelial cells and anti-Repo (blue) to identify glial cells. (D) Frontal view of volume-rendering 3D reconstructions of a wild type early L3 optic lobe showing DL1 progeny (green) in the same region as ClC-a$^+$ cells in Figure 5A. Neuroepithelia were segmented and the rest of the signal masked to avoid background noise and allow better visualization. (E) Horizontal view of a confocal plane showing the neural progeny of the DL1 lineage in the central brain (Repo$^-$) and the $Xg_o$ and $Xg_i$ glial progeny in the optic lobe (Repo$^+$). (F) Frontal view of a volume-rendering 3D reconstruction of a wild type mid L3 brain showing DL1 progeny (green) in the same region as *ClC-a*$^+$ cells in Figure 5B. (G, H) *ClC-a* lineage tracing, performed with the *05423*$^{ClC\text{-}a\text{-}GAL4}$ driver and the G-TRACE system, to analyze the drop in ClC-a$^+$ cells from mid to late L3. Anti-Repo was used to label glial cells. (G) Frontal view of a volume-rendering 3D reconstruction of a late L3 brain. (G') G-TRACE green labeling indicates that cells expressed *ClC-a* at some point during development. (G") G-TRACE red labeling shows cells currently expressing *ClC-a*. Bracket and arrowhead respectively demarcate $Xg_o$ and pag that had already expressed *ClC-a* in early and mid L3 (K'), but downregulated *ClC-a* expression in late L3 (K"). (H) Green box plots show the number of nuclei per brain that expressed *ClC-a* at a developmental time prior to the larval stage analyzed. Red box plots show the number of nuclei per brain that currently express *ClC-a*. Dots in box plots represent data points. Comparisons between green and red box plots are shown for each developmental time analyzed. *p*-values were calculated with the non-parametric Mann-Whitney test. Both in mid and late L3, there are more cells that used to express *ClC-a* than cells that currently express it, indicating that some downregulation of *ClC-a* expression is occurring at mid and late L3 stages. The number of nuclei currently expressing *ClC-a* in mid L3 quantified with *05423*$^{ClC\text{-}a\text{-}GAL4}$ mediated G-TRACE (*UAS-nsl-DsRed*) is lower than the number of cells observed when using *05423*$^{ClC\text{-}a\text{-}GAL4}$ and *UAS-H2B-RFP*, presumably due to the use of different UAS reporters. Compare mid L3 current G-TRACE value (red box plot) in (H) to the value in *05423*$^{ClC\text{-}a\text{-}GAL4}$/+ animals at mid L3 in Figure 5M. (I, J) Assessment of cell death in mutants in the region where boundary glia would normally be positioned. Early L3 (I) and mid L3 (J) mutant brains showing, as expected, few (I) or no (J) ClC-a$^+$ boundary glia nuclei (red). Most of the sporadic Dcp-1 signal observed (arrowheads, gray) is in non-glial cells (repo$^-$).

OPC, outer proliferation center; IPC, inner proliferation center; sg, satellite glia; bg, boundary glia; pag, palisade glia; $Xg_o$, outer chiasm glia; $Xg_i$, inner chiasm glia; cxg, cortex glia. Scale bars represent 10 µm. n.s.>0.05, * $p<0.05$, ** $p<0.01$.

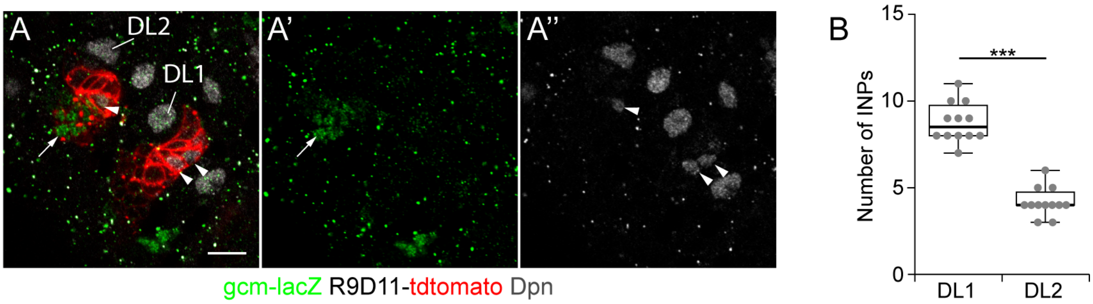


**Supplementary Figure 9. DL1/DL2 distinction based on *gcm* expression.**
(A-A") Confocal sections showing the progeny of DL1 and DL2 neuroblasts labeled by R9D11-tdtom expression (red). *gcm-lacZ* expression (green, A') labels part of the DL2 lineage (arrow). INPs (arrowheads) are labeled in anti-Deadpan antibody (gray, A"). (B) Quantification and comparison of the number of INPs per lineage. *p*-value was calculated with the non-parametric Mann-Whitney test. Scale bars represent 10 µm. ***p<0.001.

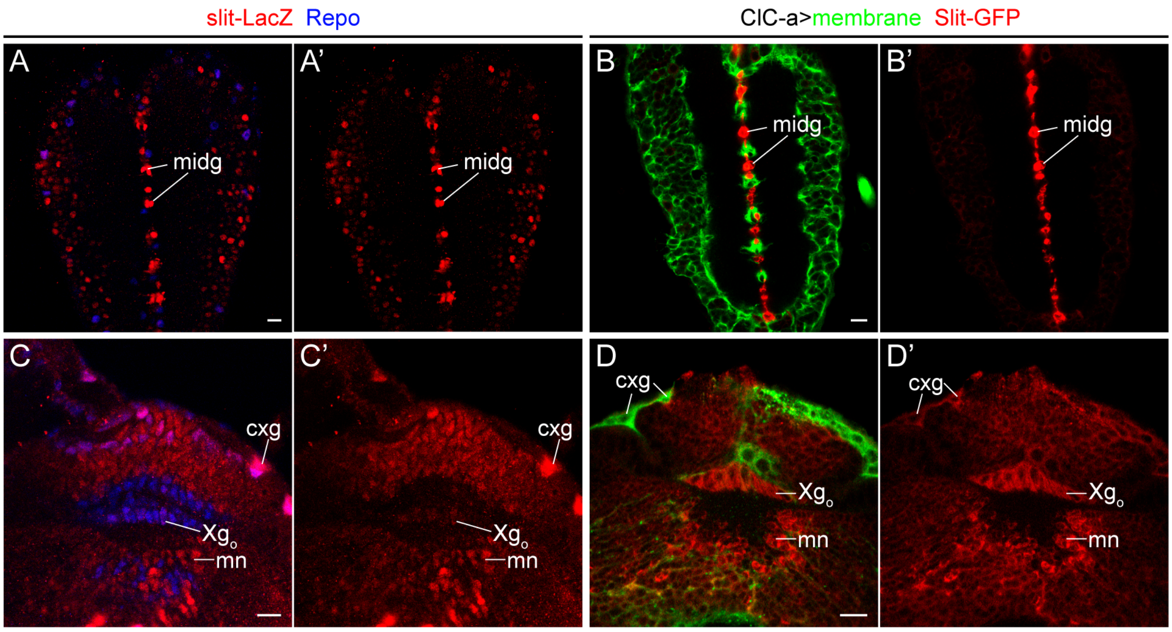


**Supplementary Figure 10. Comparison of *slit*-LacZ (*sli* $^{05428}$) and Slit-GFP (*sli[MI03825-GFSTF.2]*) expression patterns.**
*sli* $^{05428}$ is a commonly used nuclear lacZ reporter of *slit* expression. We characterized Slit-GFP expression pattern because *sli* $^{05428}$ nuclear LacZ expression at early and mid L3 stages was very low and difficult to distinguish from background. Slit full-length protein can be cleaved into large N-terminal (Slit-N) and short C-terminal (Slit-C) fragments. Slit-FL and Slit-N are more tightly associated with the cell surface, whereas Slit-C is mostly shed into the extracellular space (Brose et al., 1999). The GFP tag in this Slit-GFP reporter line was located between amino acids 398-399, in the second LRR repeat, so in the Slit-N terminal fragment. Thus, the GFP signal of the Slit-GFP reporter stays in the membrane of the *slit* expressing cells. *slit* signal for both *slit*-lacZ and Slit-GFP reporters is shown in red. *ClC-a*$^{+}$ membranes are labeled with *05423*$^{ClC\text{-}a\text{-}GAL4}$*/UAS-mCD8-mRFP* and shown in green. Glia nuclei are labeled with anti-Repo antibody (blue). (A, B) Horizontal views through the VNC showing nuclear LacZ signal (red, A, A') and membrane Slit-GFP signal (red, B, B') in midline glia. (C, D) Frontal views of late L3 optic lobes. (C) Nuclear LacZ signal (red) can be seen in Xg$_o$ and medulla neurons as previously reported (Suzuki et al., 2016; Tayler et al., 2004), as well as in cortex glia. In early pupal stages, LacZ expression in Xg$_o$ is stronger than in late L3 (data not shown). (D) Membrane Slit-GFP signal (red) is seen in the same cell types as *slit*-LacZ: Xg$_o$, cortex glia, and medulla neurons. Thus, the Slit-GFP expression pattern is the same as the one observed with *slit*-lacZ.
midg, midline glia; cxg, cortex glia; pag, palisade glia; Xg$_o$, outer chiasm glia; mn, medulla neuron. Scale bars represent 10 µm.

### GENOTYPE LIST

**Figure 1:**
**A, B, C, D, E, F, G, H, I, J.** *w; UAS-mCD8GFP/+; 05423*$^{ClC\text{-}a\text{-}GAL4}$*/UAS-H2BRFP*
**K.** *w; mir8-GAL4/UAS-mCD8GFP; +/+*
**L.** *w; UAS-mCD8GFP/+; R54H02-GAL4/UAS-H2BRFP*
**M, N.** *w; UAS-mCD8GFP, lexAop-CD2RFP/+; 05423*$^{ClC\text{-}a\text{-}GAL4}$*/wrapper932i-LexA*
**O, P, Q.** *w; UAS-mCD8GFP/+; 05423*$^{ClC\text{-}a\text{-}GAL4}$*/wrapper932i-Gal80*

**Figure 2:**
**C.**
*+/+: w*$^{1118}$*; +/+; +/+*
*05423*$^{ClC\text{-}a\text{-}GAL4}$*/+: w; +/+; 05423*$^{ClC\text{-}aGAL4}$*/+*
*14007/+: w; +/+; 14007/+*
*Df/+: w; +/+; Df(3R)PS2/+*
*05423*$^{ClC\text{-}a\text{-}GAL4}$*/14007: w; +/+; 05423*$^{ClC\text{-}aGAL4}$*/14007*
*05423*$^{ClC\text{-}a\text{-}GAL4}$*/Df: w; +/+; 05423*$^{ClC\text{-}aGAL4}$*/Df(3R)PS2*
**D.**
*UAS-ClC-a/+: w; UAS-ClC-a/+; +/+*
*UAS-ClCN2/+: w; UAS-ClCN2/+; +/+*
*05423*$^{ClC\text{-}a\text{-}GAL4}$*/Df & +/+: w; +/+; 05423*$^{ClC\text{-}a\text{-}GAL4}$*/Df(3R)PS2*
*05423*$^{ClC\text{-}a\text{-}GAL4}$*/Df & UAS-ClC-a/+: w; UAS-ClC-a/+; 05423*$^{ClC\text{-}a\text{-}GAL4}$*/Df(3R)PS2*
*05423*$^{ClC\text{-}a\text{-}GAL4}$*/Df & UAS-ClCN2/+: w; UAS-rClC-2/+; 05423*$^{ClC\text{-}a\text{-}GAL4}$*/Df(3R)PS2*
**E.**
*14007/+: w; +/+; 14007/+*
**F.**
*14007/Df: w; +/+; 14007/Df(3R)PS2*
**G.**
*05423*$^{ClC\text{-}a\text{-}GAL4}$*/+: w; +/+; 05423*$^{ClC\text{-}a\text{-}GAL4}$*/+*
*05423/+: w; +/+; 05423/+*
*14007/+: w; +/+; 14007/+*
*Df/+: w; +/+; Df(3R)PS2/+*
*05423*$^{ClC\text{-}a\text{-}GAL4}$*/Df: w; +/+; 05423*$^{ClC\text{-}a\text{-}GAL4}$*/Df(3R)PS2*
*05423/Df: w; +/+; 05423/Df(3R)PS2*
*14007/Df: w; +/+; 14007/Df(3R)PS2*
*05423*$^{ClC\text{-}a\text{-}GAL4}$*/14007: w; +/+; 05423*$^{ClC\text{-}a\text{-}GAL4}$*/14007*
*05423/14007: w; +/+; 05423/14007*
*14007/14007: w; +/+; 14007/14007*
**H.**
*Repo-GAL4/+: w; +/+; Repo-Gal4/+*
*05423*$^{ClC\text{-}a\text{-}GAL4}$*/+: w; +/+; 05423*$^{ClC\text{-}a\text{-}GAL4}$*/+*
*UAS-ClC-aRNAi/+: w; UAS-ClC-aRNAi/+; UAS-Dcr2/+*
*Repo>ClC-aRNAi: w; UAS-ClC-aRNAi /+; Repo-GAL4/UAS-Dcr2*
*05423*$^{ClC\text{-}a\text{-}GAL4}$*>ClC-aRNAi: w; UAS-ClC-aRNAi /+; 05423*$^{ClC\text{-}a\text{-}GAL4}$*/UAS-Dcr2*
**I.**
*Repo-GAL4/+ & 14007/Df(3R)PS2: w; Repo-GAL4/+; 14007/Df(3R)PS2*
*UAS-ClC-a/+ & 14007/Df(3R)PS2: w; UAS-ClC-a/+; 14007/Df(3R)PS2*
*UAS-ClCN2/+ & 14007/Df(3R)PS2: w; UAS-ClCN2/+; 14007/Df(3R)PS2*
*Repo>ClC-a & 14007/Df(3R)PS2: w; UAS-ClC-a/Repo-GAL4; 14007/Df(3R)PS2*
*Repo>ClCN2 & 14007/Df(3R)PS2: w; UAS-ClCN2/Repo-GAL4; 14007/Df(3R)PS2*

*05423$^{ClC-a-GAL4}$/Df & +/+: w; +/+; 05423$^{ClC-a-GAL4}$/Df(3R)PS2*
*05423$^{ClC-a-GAL4}$/Df & UAS-ClC-a/+: w; UAS-ClC-a/+; 05423$^{ClC-a-GAL4}$/Df(3R)PS2*
*05423$^{ClC-a-GAL4}$/Df & UAS-ClCN2/+: w; UAS-rClC-2/+; 05423$^{ClC-a-GAL4}$/Df(3R)PS2*

**Figure 3:**
**A, B, C, D.**
*05423$^{ClC-a-GAL4}$/+: w; UAS-mCD8GFP/+; 05423$^{ClC-a-GAL4}$/UAS-H2BRFP*
**E, F, G, H.**
*05423$^{ClC-a-GAL4}$/14007: w; UAS-mCD8RFP/UAS-H2BYFP; 05423$^{ClC-aGAL4}$/14007*

**Figure 4:**
**A, B, C, F, G.**
*14007/+: w; +/+; 14007/+*
*14007/Df: w; +/+; 14007/Df(3R)PS2*
**D, E.**
*14007/+: hsFLP, FRT19A, tub-Gal80/FRT19A; tub-GAL4,UAS-mCD8GFP /+; 14007/+*
*14007/Df: hsFLP, FRT19A, tub-Gal80/FRT19A; tub-GAL4,UAS-mCD8GFP /+; 14007/Df(3R)PS2*
**H, I.**
*+/+: w; +/+; +/+*
*14007/+: w; +/+; 14007/+*
*Df/+: w; +/+; Df(3R)PS2/+*
*14007/Df: w; +/+; 14007/Df(3R)PS2*
**J, K.**
*mir8$^{cxg}$ or UAS-ClC-a control: tub>Gal80>/+; mir8-GAL4/+; 14007/Df(3R)PS2 or tub>Gal80>/+; +/RepoFLP6.2, UAS-ClC-a; 14007/Df(3R)PS2*
*mir8$^{cxg}$>ClC-a: tub>Gal80>/+; mir8-GAL4/RepoFLP6.2, UAS-ClC-a; 14007/Df(3R)PS2*

**Figure 5:**
**A, B, C, D, E, F.**
*05423$^{ClC-a-GAL4}$/+: w; UAS-mCD8GFP/+; 05423$^{ClC-a-GAL4}$/UAS-H2BRFP*
**G, H, I, J, K, L.**
*05423$^{ClC-a-GAL4}$/14007: w; UAS-mCD8RFP/UAS-H2BYFP; 05423$^{ClC-aGAL4}$/14007*
**M.**
*05423$^{ClC-a-GAL4}$/+: w; UAS-mCD8RFP/ UAS-H2BYFP; 05423$^{ClC-aGAL4}$/+*
*05423$^{ClC-a-GAL4}$/14007: w; UAS-mCD8RFP/ UAS-H2BYFP; 05423$^{ClC-aGAL4}$/14007*
**O.**
*14007/+: w; +/+; 14007, R9D11-tdtomato/+*
**P, Q.**
*14007/Df: w; +/+; 14007, R9D11-tdtomato/Df(3R)PS2*
**R, S, U.**
*05423$^{ClC-a-GAL4}$/+: w; UASmCD8GFP/+; 05423$^{ClC-aGAL4}$/R9D11-tdtomato*
**T,U.**
*05423$^{ClC-a-GAL4}$/14007: w; UASmCD8GFP/+; 05423$^{ClC-aGAL4}$/14007, R9D11-tdtomato*

**Figure 6:**
**A, B, C, D.**
*05423$^{ClC-a-GAL4}$/+: w; UAS-mCD8GFP/+; 05423$^{ClC-a-GAL4}$/UAS-H2BRFP*
**E, F.**
*14007/Df: w; +/+; 14007/Df(3R)PS2*

**G, H.**

slit$^{dui}$ / slit$^{dui}$: w; slit$^{dui}$, GMR-GFP/slit$^{dui}$, GMR-GFP; +/+

**I, J, K.** w; UAS-mCD8RFP/Slit-GFP; 05423$^{ClC\text{-}a\text{-}GAL4}$/+

**L.**

14007/+: w; +/+; 14007/+

slit$^{dui}$ /+: w; slit$^{dui}$, GMR-GFP/+; 14007/+

14007/14007: w; +/+; 14007/14007

slit$^{dui}$ /+; 14007/14007: w; slit$^{dui}$, GMR-GFP/+; 14007/14007

**M.**

05423$^{ClC\text{-}a\text{-}GAL4}$/+: w; +/+; 05423$^{ClC\text{-}a\text{-}GAL4}$/+

UAS-slitRNAi/+: w; UAS-slitRNAi/+; UAS-Dcr2/+

05423$^{ClC\text{-}a\text{-}GAL4}$>slitRNAi: w; UAS-slitRNAi/+; UAS-Dcr2/05423$^{ClC\text{-}a\text{-}GAL4}$

slit$^{dui}$/+; 05423$^{ClC\text{-}a\text{-}GAL4}$>slitRNAi: w; UAS-SlitRNAi/ slit$^{dui}$, GMR-GFP; UAS-Dcr2/05423$^{ClC\text{-}a\text{-}GAL4}$

**Figure 7:**

**B.**

14007/+: w; +/+; 14007/+

14007/Df: w; +/+; 14007/Df(3R)PS2

14007/Df, mir8$^{cxg}$>ClC-a: tub>Gal80>/+; mir8-GAL4/RepoFLP6:2, UAS-ClC-a;14007/Df(3R)PS2

**C.**

mir8$^{cxg}$ or UAS-ClC-a control: tub>Gal80>/+; mir8-GAL4/+; 14007/Df(3R)PS2 or tub>Gal80>/+; +/RepoFLP6.2, UAS-ClC-a; 14007/Df(3R)PS2

mir8$^{cxg}$>ClC-a: tub>Gal80>/+; mir8-GAL4/RepoFLP6.2, UASClC-a; 14007/Df(3R)PS2

**Figure 8:**

**B, D.**

05423$^{ClC\text{-}a\text{-}GAL4}$/+: w; UASmCD8GFP/+; 05423$^{ClC\text{-}aGAL4}$/+

**C.**

05423$^{ClC\text{-}a\text{-}GAL4}$/+: w; +/+; 05423$^{ClC\text{-}aGAL4}$/+

**E.**

14007/+: w; +/+; 14007/+

**F, H.**

05423$^{ClC\text{-}a\text{-}GAL4}$/14007: w; UASmCD8GFP/+; 05423$^{ClC\text{-}aGAL4}$/14007

**G.**

05423$^{ClC\text{-}a\text{-}GAL4}$/14007: w; +/+; 05423$^{ClC\text{-}aGAL4}$/14007

**I.**

14007/Df: w; +/+; 14007/Df(3R)PS2

**K, L.**

14007/+: hsFLP,FRT19A,tubGal80/FRT19A; tubGAL4,UASmCD8GFP/+; 14007/+

**M, N, O.**

14007/Df: hsFLP,FRT19A,tubGal80/FRT19A; tubGAL4,UASmCD8GFP/+; 14007/Df(3R)PS2

**Supplementary figure 1:**

**A, D, G.** w; +/+; +/+

**B, E, H.** w; +/+; ClC-aGFP/ClC-aGFP

**C, F.** w; UAS-mCD8GFP/+; 05423$^{ClC\text{-}aGAL4}$/+

**I.** w; UAS-mCD8GFP/+; 05423$^{ClC\text{-}aGAL4}$/UAS-H2BRFP

**Supplementary figure 2:**

**A, B, C, D, E, F, G.** w; UAS-mCD8GFP/+; 05423$^{ClC\text{-}aGAL4}$/UAS-H2BRFP

**Supplementary figure 3:**
**A, C.**
*+/+: w; +/+; +/+*
**B, D.**
*14007/Df: w; +/+; 14007/Df(3R)PS2*
**E.**
*+/+: w; +/+; +/+*
*Df/+: w; +/+; Df(3R)PS2/+*
*05423/+: w; +/+; 05423/+*
*14007/+: w; +/+; 14007/+*
*05423/Df: w; +/+; 05423/Df(3R)PS2*
*14007/Df: w; +/+; 14007/Df(3R)PS2*
*14007/05423: w; +/+; 14007/05423*

**Supplementary figure 4:**
**A.** *w; Rh1-GAL4/UAS-mCD8GFP; 14007/+*
**B, C.** *w; Rh1-GAL4/UAS-mCD8GFP; 14007/Df(3R)PS2*
**D.** *w; Rh4-EGFP/+; 14007/+*
**E.** *w; Rh4-EGFP/+; 14007/Df(3R)PS2*
**F.** *w; Rh6-LacZ/+; 14007/+*
**G.** *w; Rh6-LacZ/+; 14007/Df(3R)PS2*
**H.** *w; sens-GAL4, UAS-utrGFP/+; 14007/+*
**I.** *w; sens-GAL4, UAS-utrGFP /+; 14007/Df(3R)PS2*
**J.** *w; Rh6-LacZ/ Rh4-EGFP; 14007/Df(3R)PS2*
**K.**
*GMR-GAL4/+: w; GMR-GAL4/+; +/+*
*UAS-ClC-aRNAi/+: w; UAS-ClC-aRNAi/+; UAS-Dcr2/+*
*GMR>ClC-aRNAi: w; UAS-ClC-aRNAi/GMR-GAL4; UAS-Dcr2/+*
*GMR-GAL4/+, 14007/Df: w; GMR-GAL4/+; 14007/Df(3R)PS2*
*UAS-ClC-a/+, 14007/Df: w; UAS-ClC-a/+; 14007/Df(3R)PS2*
*GMR>ClC-a, 14007/Df: w; GMR-GAL4/UAS-ClC-a; 14007/Df(3R)PS2*

**Supplementary figure 5:**
**A.**
$05423^{ClC\text{-}a\text{-}GAL4}$*/+: w; UAS-mCD8GFP/ UAS-H2BYFP;* $05423^{ClC\text{-}a\text{-}GAL4}$*/+*
$05423^{ClC\text{-}a\text{-}GAL4}$*/14007: w; UAS-mCD8RFP/ UAS-H2BYFP;* $05423^{ClC\text{-}a\text{-}GAL4}$*/14007*
**B.**
*14007/+: w; +/+; 14007/+*
*14007/Df: w; +/+; 14007/Df(3R)PS2*

**Supplementary figure 6:**
**A, D.**
*14007/+: w; +/+; 14007/+*
**B, C, E.**
*14007/Df: w; +/+; 14007/Df(3R)PS2*

**Supplementary figure 7:**
**A, C.** *w; mir8-GAL4/UAS-mCD8GFP; +/+*
**B, D, E.** *tub>GAL80>/+; mir8-GAL4, RepoFLP6.2/+; UAS-mCD8GFP/+*

**Supplementary figure 8:**
**A, B.** *w; UAS-mCD8GFP, lexAop-CD2RFP/R43H01-LexA; 05423*$^{ClC\text{-}a\text{-}GAL4}$*/+*
**C.** *w; UAS-mCD8GFP/+; R25A01-GAL4/+*
**D, E, F.** *w; UAS-G-TRACE/+; R38H02-GAL4/+*
**G, H.** *w; UAS-G-TRACE/+; 05423*$^{ClC\text{-}a\text{-}GAL4}$*/+*
**I, J**
*05423*$^{ClC\text{-}a\text{-}GAL4}$*/14007:* *w; UAS-H2BYFP/+; 05423*$^{ClC\text{-}a\text{-}GAL4}$*/14007*

**Supplementary figure 9:**
**A.** *w; gcm-lacZ/+; R9D11-tdtomato/+*

**Supplementary figure 10:**
**A, C.** *w; slit-lacZ/+; +/+*
**B, D.** *w; slit-GFP/UAS-mCD8RFP; 05423*$^{ClC\text{-}a\text{-}GAL4}$*/+*
